# WGA-LP: a pipeline for Whole Genome Assembly of contaminated reads

**DOI:** 10.1101/2021.07.31.454518

**Authors:** N. Rossi, A. Colautti, L. Iacumin, C. Piazza

## Abstract

**Summary:** Whole Genome Assembly (WGA) of bacterial genomes with short reads is a quite common task as DNA sequencing has become cheaper with the advances of its technology. The process of assembling a genome has no absolute golden standard (Del Angel et al. (2018)) and it requires to perform a sequence of steps each of which can involve combinations of many different tools. However, the quality of the final assembly is always strongly related to the quality of the input data. With this in mind we built WGA-LP, a package that connects state-of-art programs and novel scripts to check and improve the quality of both samples and resulting assemblies.

WGA-LP, with its conservative decontamination approach, has shown to be capable of creating high quality assemblies even in the case of contaminated reads.

**Availability and Implementation:** WGA-LP is available on GitHub (https://github.com/redsnic/WGA-LP) and Docker Hub (https://hub.docker.com/r/redsnic/wgalp). The web app for node visualization is hosted by shinyapps.io (https://redsnic.shinyapps.io/ContigCoverageVisualizer/).

**Contact:** Nicolò Rossi

**Supplementary information:** Supplementary data are available at *bioRxiv* online.

## 1 Introduction

A currently active challenge in the context of whole genome assembly for bacterial genomes is to produce reliable whole genome assemblies that are contaminant free (Steinegger and Salzberg (2020) and Chun et al. (2018)). In this context, we built WGA-LP, a pipeline that includes different strategies to guide the users in producing higher quality whole genome assemblies, by also including specific features to control possible contamination. Moreover, its workflow is structured to assist in the quality evaluation of the results of each step of the pipeline by providing useful plots and summaries. The current state-of-the-art for decontamination consists in the use of Kraken2, (Wood et al. (2019)) a software for read origin imputation, and of pipelines like ProDeGe (Tennessen et al. (2016)) and SIDR (Fierst and Murdock (2017)). This last is however meant for eukaryotic genomes.

## 2 Software description

WGA-LP software is built to be used from the command line. The procedures of the pipeline are organized by functionality and have a consistent syntax for argument passing. More details are available in the Supplementary Information material, on the GitHub, and Docker Hub web pages of the tool.

WGA-LP performs many steps that can be run independently. In order to execute the whole workflow the user is required to provide the raw reads (`.fastq`) and, optionally, the references that should be used for decontamination (`.fasta`). All the other input files can be produced using WGA-LP commands. Check the Supplementary information material for a complete explanation of all the input parameters for WGA-LP.

The first step of WGA-LP has the role of assessing the quality of the input reads and detecting possible contamination sources. To this end WGA-LP relies on Trimmomatic (Bolger et al. (2014)), FastQC (Andrews (2010)), Kraken2, and Bracken (Lu et al. (2017)). The trimming step is fully configurable so that the user can choose the right approach for his data.

The decontamination procedure is based on a custom script that includes calls to three programs: BWA mem (Li (2013)), Samtools, (Li et al. (2009)) and Bazam (Sadedin and Oshlack (2019)). We take in input the raw reads and two sets of references, one for the target organism and one for the contaminants. This procedure is presented with more details in the Figure 1 and in the Supplementary materials. This decontamination approach is conservative and reduces the probability of discarding reads of the target organism. This part of the pipeline can be used as a standalone program and can be combined with any other program for Whole Genome Assembly.

**Figure 1:**
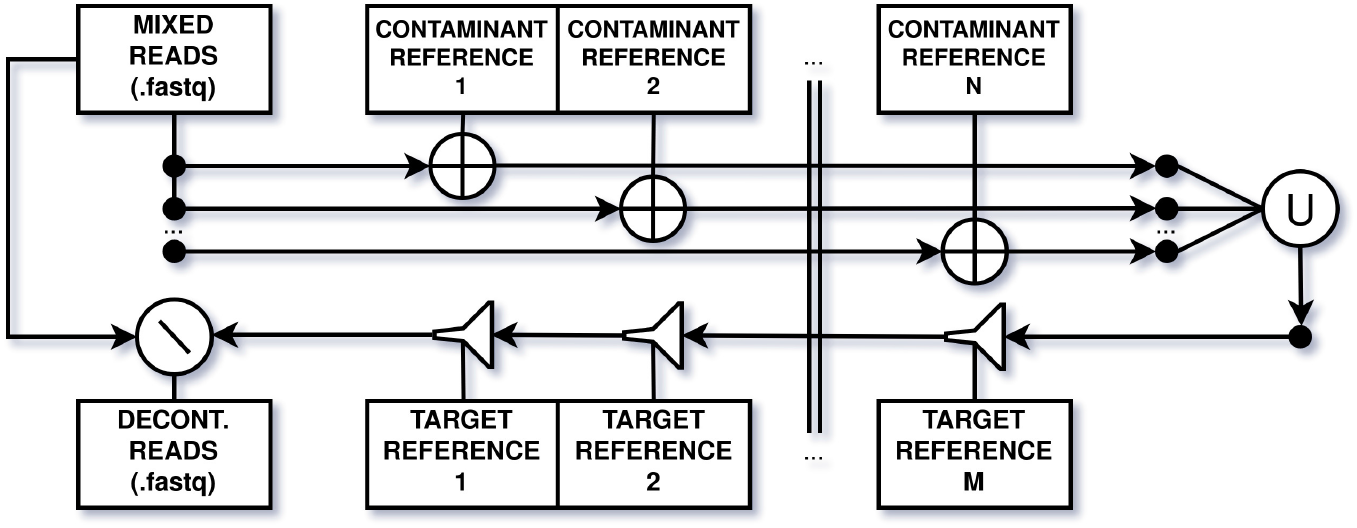
The decontamination procedure. Input reads are mapped against each reference of the contaminant independently (first three wires from left to are then merged together (Union, ∪) and gradually filtered (last wire from right to left), with the effect of removing all the reads that map to any The final decontaminated reads are extracted by set difference (\) using the original input set.

WGA-LP natively supports SPAdes (Bankevich et al. (2012)) and Minia (Chikhi and Rizk (2012)) assemblers. SPAdes is currently a common choice for bacterial WGA, while Minia is a very simple and fast assembler. The other steps of WGA-LP can support any assembler that includes in its outputs a fasta formatted assembly and a fastg assembly graph (required only for putative plasmid search with Recycler (Rozov et al. (2017))).

WGA-LP includes custom scripts to help in the visualization of node coverage by post processing the output of Samtools depth. This allows to produce coverage plots (computed by remapping the reads to the assembled genome) that can be helpful in finding anomalies, such as prophage insertions in the genome. Moreover, WGA-LP provides a web app and tools for nodes (and reads) selection that can improve the decontamination results. These act by exploiting the assembly process as it tends to assemble nodes with reads of the same organism. Such procedures are well fitted to be combined with Kraken2, since this tool can point out problematic nodes, that can be then further evaluated with BLAST alignment (Altschul et al. (1990)) in order to validate user selections.

For node reordering, WGA-LP uses the `ContigOrderer` option from Mauve aligner (Rissman et al. (2009)). This step requires to provide a reference for the target organism.

WGA-LP offers interfaces to two programs that extract putative plasmids: plasmidSPAdes (Antipov et al. (2016)) and Recycler. It is highly recommended to check the results of these tools using BLAST.

WGA-LP includes three programs to evaluate the quality of the final result of the pipeline: Quast (Gurevich et al. (2013)) CheckM (Parks et al. (2015)) and Merqury (Rhie et al. (2020)). Especially, checkM is useful to verify the completeness and contamination of the produced assembly.

For the annotation, WGA-LP interfaces with Prokka (Seemann (2014)) in order to create NCBI compliant assemblies. This can be considered the final output of the pipeline and can be used for downstream analysis.

## 3 Results

We tested WGA-LP pipeline on real data (see Data Availability section) and we have shown how its workflow was effective in producing an high quality Whole Genome Assembly even in this challenging scenario of a contaminated genome, with improvements in comparison with less curated approaches (see Supplementary Information). Finally, we extended the comparison to include ProDeGe, another state-of-the-art decontamination procedure. ProDeGe alone was not able to filter large nodes of the contaminant, however it was possible to use WGA-LP procedures based on kraken2 classification to refine the resulting assembly, achieving comparable results with our pipeline. However, also in this case our tool performed better on the elimination of the shorter nodes, keeping those that, in a further check, were classified from the target genome by BLAST alignment.

Both the decontamination procedure and the node selection that are the core of our pipeline, can be integrated in any other pipeline for Whole Genome Assembly, in the pre-processing and post-processing phases.

## 4 Data availability

The testing reads, from the organism Lacticaseibacillus rhamnosus, heavily contaminated with Pediococcus acidilactici, were deposited in the NCBI’s Sequence Read Archive (SRA) with the ID `SRR15265000`, included in the associated BioProject `PRJNA749304`.

## Acknowledgements

All website and links in this paper and in the supplementary materials were accessed on the 25/07/2021.

## Funding

This work was funded by Ministero dell’Università e della Ricerca, Rome, Italy, FIRB n. RBFR107VML.

## Notes

https://github.com/redsnic/WGA-LP

https://hub.docker.com/r/redsnic/wgalp

https://redsnic.shinyapps.io/ContigCoverageVisualizer/

### Competing Interest Statement

The authors have declared no competing interest.

### Summary of Updates

This article has been accepted for publication in Bioinformatics Published by Oxford University Press.

