## Supplementary Material for "WGA-LP: a pipeline for Whole Genome Assembly of contaminated reads" (from another version of this preprint: version 1, 2021-08-03)

Nicolò Rossi, Andrea Colautti, Lucilla Iacumin, and Carla Piazza

24-07-2021

### Supplementary material

#### Setting up WGA-LP

##### Docker Installation

> This steps requires to have `Docker` installed on your machine. Guides to install it are available at the website https://docs.docker.com/get-docker/.

Using `Docker` installation allows to:

- manage all the dependencies of WGA-LP automatically
- create a controlled environment for the analysis
- include all the databases needed to make the tools of WGA-LP work
- choose a preferred operating system, as `Docker` interfaces are available for `Linux`, `MacOS`, and `Windows`.

Installing WGA-LP from docker is quite straightforward:

```
# pull image
docker pull redsnic/wgalp:1.00
# run image: replace <HOST_SHARED_FOLDER> with a directory on your file system
docker run -itd -v <HOST_SHARED_FOLDER>:/root/shared --name wgalp
↪  redsnic/wgalp:1.00
# access shell
docker exec -it wgalp /bin/bash
```

**Optional**: To be able to create RAMdisks for the kraken2 database, a privileged container is needed. Replace the previous `docker run` command with the following:

```
docker run -itd -v <HOST_SHARED_FOLDER>:/root/shared --privileged --name wgalp
↪  redsnic/wgalp:1.00
```

> in practice, this affect only the commands `wgalp kdb-load` and `wgalp kbd-unload`. Note that the host will need root permission to access folders and files created by privileged containers.

The execution of these command should print out a welcome message and show the WGA-LP custom prompt:

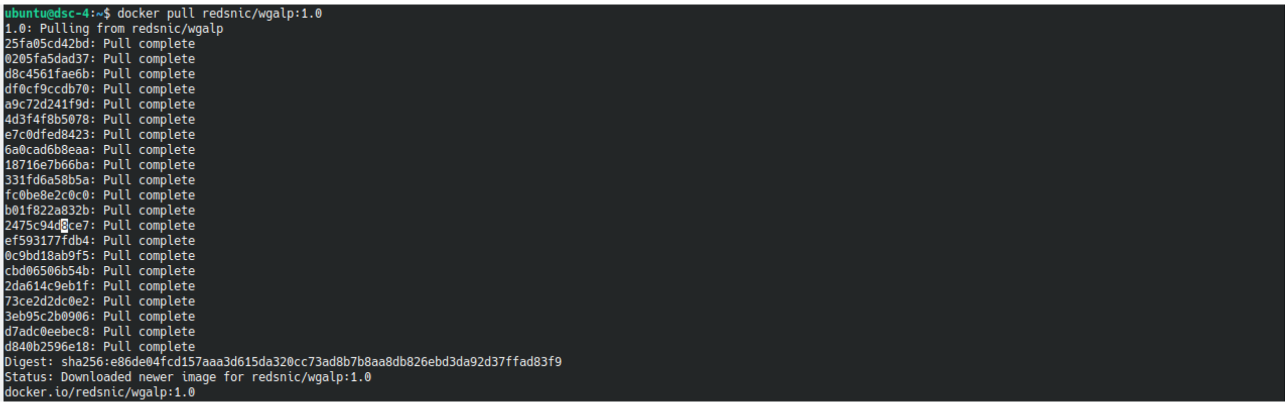

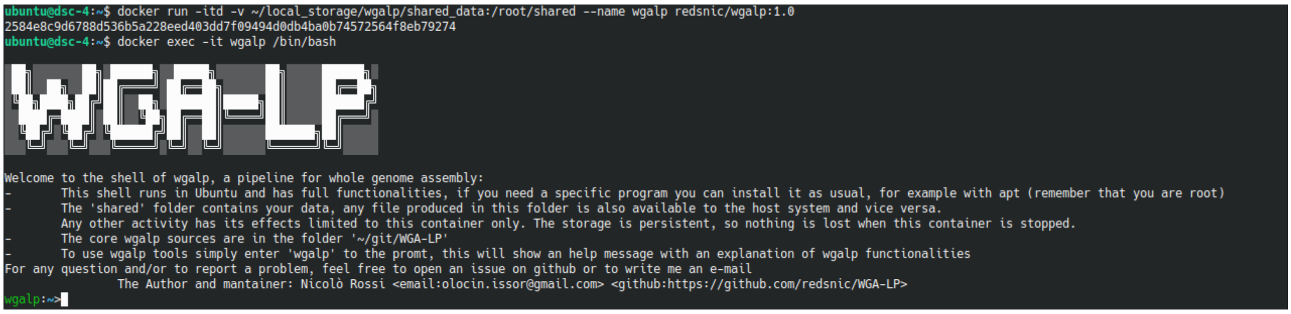

When inside the WGA-LP shell, a full `Ubuntu 18.04` environment is available, so the user has all the `bash` functionalities, including package managers like `conda` and `apt`.

> The user is `root` in the WGA-LP shell. For this reason, avoid using the `sudo` command as it is not required and will generate an error. Note that these permission are limited to the container space, as usual with `Docker`.

You can check WGA-LP's help message by simply typing `wgalp` to the prompt:

```
wgalp:~>wgalp
 --- wgalp: a pipeline for bacterial Whole Genome Assembly ---

This programs is an helper to run the sub procedures of wgalp
usage: wgalp <program> [args]

the following is a list of all the available programs:
        trim : trim reads and/or assess contaminations with kraken2
        decontaminate : remove reads mapping to a contaminant non ambiguosly
        assemble : assemble reads into scaffolds or contigs
        check-coverage : compute coverage statistics of an assembled genome
        view-nodes : compute coverage plots for specific nodes of a whole genome assembly       reorder : reorder a whole genome assembly using a reference genome
        filter-assembly : select contigs by ID
                          (to be used with the webapp: https://redsnic.shinyapps.io/ContigCoverageVisualizer/)
        annotate : run prokka annotation with NCBI standard
        plasmid : extract putative plasmids using recycler
        quality : evaluate assembly quality using checkM, merqury and quast
        understand-origin : runs kraken2 in selection mode
        kdb-load : pre-load kraken2 database in RAM, so that you dont have to load it multiple times (use --memory-mapped option when possible)
        kdb-unload : remove loaded kraken2 db from RAM
        filter-fastq : select reads from a fastq file
        help : show this message (equivalent to --help or -h)

Run wgalp <program> --help for specific information about the selected program.

wgalp:~>
```

Now WGA-LP is installed on your computer and ready for use.

###### Manual installation (alternative)

Even if `Docker` installation should be preferred, it is possible to manually install WGA-LP an all its dependencies. The `Dockerfile` in the WGA-LP GitHub repository can serve as a reference guide. Remember to give `root` privileges when needed.

###### Updating WGA-LP

If you want to update WGA-LP source code to its latest version, you can do that by issuing the following commands:

```
cd /root/git/WGA-LP/ && \
    git pull && \
    pip install .
```

as it can be seen in this example:

```
wgalp:~>cd git/WGA-LP/
wgalp:~/git/WGA-LP>git pull
remote: Enumerating objects: 63, done.
remote: Counting objects: 100% (63/63), done.
remote: Compressing objects: 100% (30/30), done.
remote: Total 49 (delta 27), reused 41 (delta 19), pack-reused 0
Unpacking objects: 100% (49/49), done.
From https://github.com/redsnic/WGA-LP
   6bc51fa..aa36f60  main       -> origin/main
Updating 6bc51fa..aa36f60
Fast-forward
 Dockerfile                                          |    26 +-
 README.md                                           |   152 +-
 RScripts/eggnog_management.Rmd                      |    64 +
 RScripts/eggnog_management.nb.html                  |  1995 +++
 WGALP/blocks/FastQC.py                              |     4 +-
 WGALP/blocks/checkM.py                              |     4 +-
 WGALP/blocks/minia.py                               |     4 +-
 WGALP/blocks/samtools_stats.py                      |     2 +
 docker_bashrc.sh                                    |   118 +
 mini_programs/understand_origin.py                  |     2 +-
 other_scripts/eggnog_letterfreq.R                   |    62 +
 other_scripts/extract_gene_ids_by_presence_absence_roary.R |    16 +
 other_scripts/filter_pangenome.sh                   |    15 +
 other_scripts/get_kegg.R                            |    14 +
 other_scripts/test/filtered_contigs.fasta           | 47684 ++++++++++++++++++++++++++++++++++++++++++++++++++++++++++++++++
 wgalp.py                                            |     2 +-
 16 files changed, 50060 insertions(+), 104 deletions(-)
 create mode 100644 RScripts/eggnog_management.Rmd
 create mode 100644 RScripts/eggnog_management.nb.html
 create mode 100644 docker_bashrc.sh
 create mode 100644 other_scripts/eggnog_letterfreq.R
 create mode 100644 other_scripts/extract_gene_ids_by_presence_absence_roary.R
 create mode 100644 other_scripts/filter_pangenome.sh
 create mode 100644 other_scripts/get_kegg.R
 create mode 100644 other_scripts/test/filtered_contigs.fasta
wgalp:~/git/WGA-LP>pip install .
Processing /root/git/WGA-LP
Requirement already satisfied: wheel in /root/miniconda/lib/python3.8/site-packages (from WGALP==0.99) (0.35.1)
Requirement already satisfied: pandas in /root/miniconda/lib/python3.8/site-packages (from WGALP==0.99) (1.2.5)
Requirement already satisfied: python-dateutil>=2.7.3 in /root/miniconda/lib/python3.8/site-packages (from pandas->WGALP==0.99) (2.8.1)
Requirement already satisfied: numpy>=1.16.5 in /root/miniconda/lib/python3.8/site-packages (from pandas->WGALP==0.99) (1.21.0)
Requirement already satisfied: pytz>=2017.3 in /root/miniconda/lib/python3.8/site-packages (from pandas->WGALP==0.99) (2021.1)
Requirement already satisfied: six>=1.5 in /root/miniconda/lib/python3.8/site-packages (from python-dateutil>=2.7.3->pandas->WGALP==0.99) (1.15.0)
Building wheels for collected packages: WGALP
  Building wheel for WGALP (setup.py) ... done
  Created wheel for WGALP: filename=WGALP-0.99-py3-none-any.whl size=51204 sha256=a47b605eaef5384448b8d9b41aeae39ee16147ced8b32f2ab001c3ea5967fe27
  Stored in directory: /root/.cache/pip/wheels/f2/90/79/74b98bb86042b1c7417f3134f589927b1c36f7081c7566001c
Successfully built WGALP
Installing collected packages: WGALP
  Attempting uninstall: WGALP
    Found existing installation: WGALP 0.99
    Uninstalling WGALP-0.99:
      Successfully uninstalled WGALP-0.99
Successfully installed WGALP-0.99
wgalp:~/git/WGA-LP>
```

WGA-LP executables are already linked to the PATH at the creation of the container.

##### Testing machine and hardware requirements

The testing machine is based on an `Intel Xeon E3-12xx v2 (Ivy Bridge, IBRS)` CPU with 32 threads and 128GB of RAM. The minimum requirement to run all steps of the analysis is to have 16GB of RAM (14GB of which free). WGA-LP is, in fact, fit to run also on average consumer laptops and it has been widely tested with success.

#### Using WGA-LP

To show the how to use WGA-LP, we will present a full analysis on real data that is available on the Sequence Read Archive, within the BioProject **PRJNA749304**.

> It is recommended to run the pipeline in the `/root/shared` folder or its subfolder, so that the data is available also on the host machine.

In this example, folder `/root/shared/144` contains the raw paired end Illumina reads and folder `/root/shared/144_working_directory` will contain the results of the analysis.

> In order to avoid loss of data, the substeps of `wgalp` command do not rewrite already present results. Depending on the error, it may be necessary to delete the output folder of a failed substeps before trying again.

##### Trimming reads

To run read **trim** and **adapter removal** we can use the `wgalp trim` command:

```
(base) wgalp:~/shared/144_working_directory>wgalp trim
--- no arguments given, printing help message ---
Run quality evaluation of raw reads, prepare trimmed reads and assess contaminations
--- arguments:
        --fastq-fwd : raw forward reads (.fastq)
        --fastq-rev : raw reverse reads (.fastq)
        --output : path to the output folder
        --kraken-db : path to the (mini)kraken database
        --memory-mapped : add this flag if you don't want to load the (mini)kraken db in RAM (so, when using a ramdisk)
        --just-kraken : add this flag if you just want to run kraken/bracken (if you have, decontamined reads for example)
        --trimmomatic-args : override default trimmomatic settings, write this field as a single string
                care that this code will be interpreted direcly by the bash shell, adding ;, |, &&, etc.. may break the execution
                by default we use: SLIDINGWINDOW:5:20 ILLUMINACLIP:TruSeq2-PE.fa:2:30:10
        --help : print this message

(base) wgalp:~/shared/144_working_directory>
```

This procedure also runs **kraken2** and **bracken** to assess possible contamination in the reads.

We issue the following command:

```
wgalp trim \
    --fastq-fwd ../144/144_S13_L001_R1_001.fastq \
    --fastq-rev ../144/144_S13_L001_R2_001.fastq \
    --kraken-db $kraken_db \
    --output trimming_step
```

> For convenience, it is possible to get the location of the kraken2 database just by using the **kraken_db** environment variable. This is done just as in the previous example; the variable is defined at the setup of WGA-LP.

With the following output:

```
          >> Reads not distributed (eg. no species above threshold): 428
          >> Unclassified reads: 75255
BRACKEN OUTPUT PRODUCED: trimming_step/bracken/bracken.log
PROGRAM END TIME: 07-12-2021 09:15:19
  Bracken complete.
INFO: With outputs {'bracken_report': 'bracken.report', 'bracken_log': 'bracken.log'}
task completed successfully
trimmed reads are at the following locations:
        trimmed_fwd : trimming_step/TrimmomaticPE/144_S13_L001_R1_001.trimmed.fastq
        trimmed_rev : trimming_step/TrimmomaticPE/144_S13_L001_R2_001.trimmed.fastq
check trimming_step for kraken and bracken outputs

real    3m33.345s
user    4m7.955s
sys     0m27.579s
(base) wgalp:~/shared/144_working_directory>ls trimming_step/
TrimmomaticPE  bracken  fastqc_raw_fwd  fastqc_raw_rev  fastqc_trimmed_fwd  fastqc_trimmed_rev  kraken
(base) wgalp:~/shared/144_working_directory>
```

Notice that the main output files of the step are written with their full path at the end of the step, to make it easier to use them in the further phases of the analysis. Execution times are based on the testing machine described in the previous sections.

WGA-LP offers many quality control features. In this step it is possible to check the quality of the reads before and after trimming through the automatically generated `fastqc` reports. The following plots show the per base sequence quality in the context of our example.

Reverse reads **before** trimming:

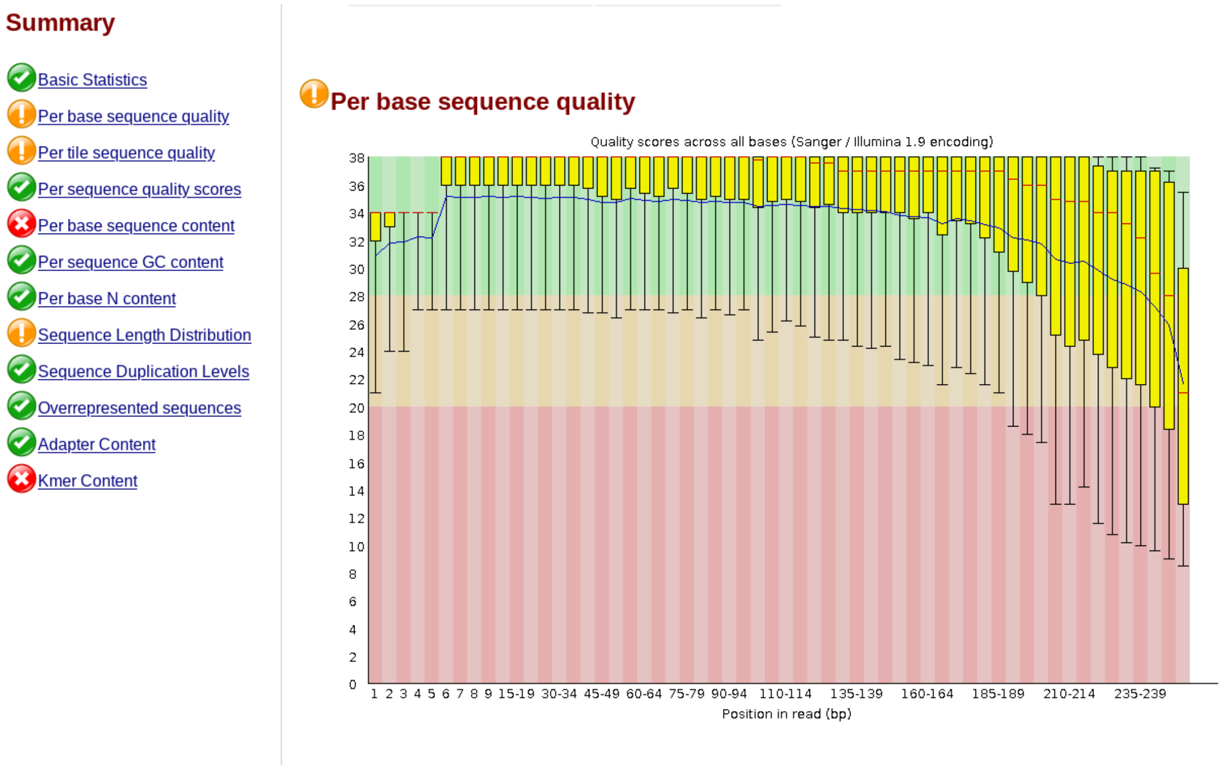

Reverse reads **after** trimming:

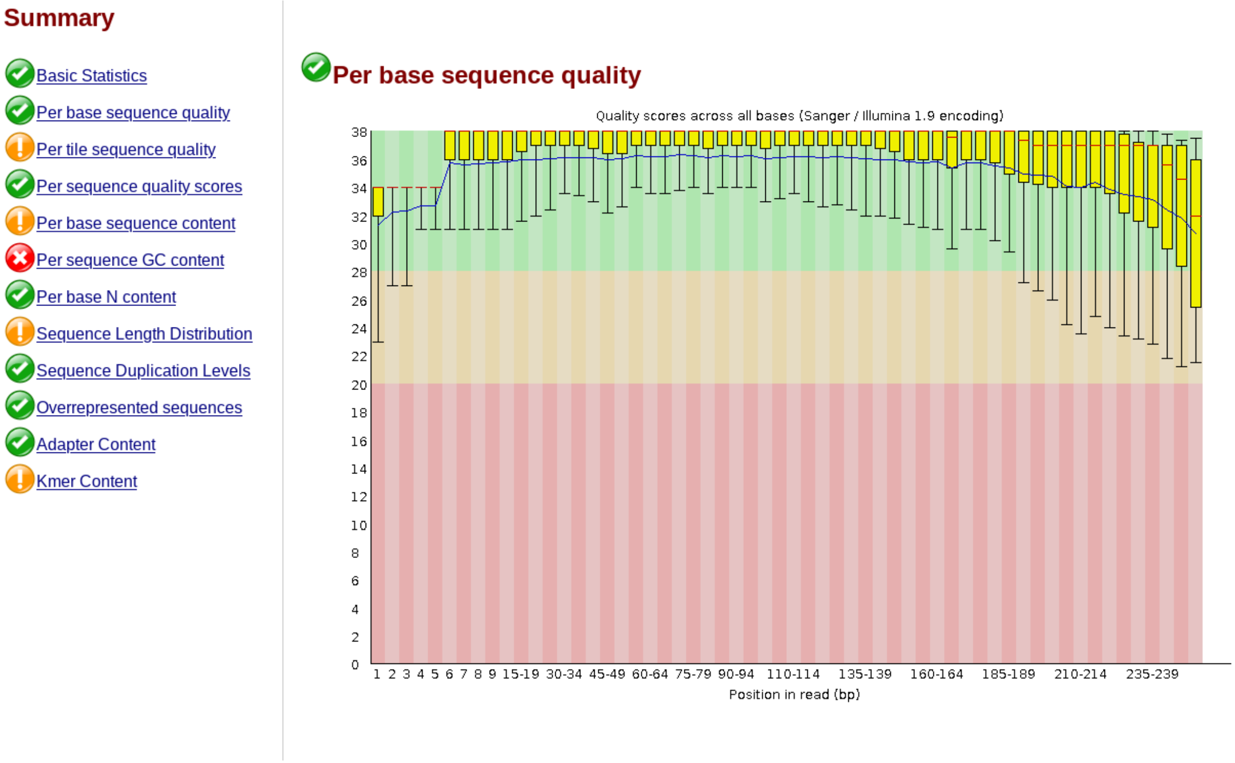

> `fastqc` reports for the reads before and after trimming must be analyzed in depth to check the quality of the results.

From the `kraken/kraken.report` file we can see that the majority of the reads map to Lactobacillus, while there is however a discrete contamination of Pediococcus:

```
...
62.94   541160   24425    G        1578                    Lactobacillus
```

```
...
7.29  62642   312     G       1253                    Pediococcus
...
```

The columns in this file must be interpreted as by kraken2 manual:

```
1. Percentage of fragments covered by the clade rooted at this taxon
2. Number of fragments covered by the clade rooted at this taxon
3. Number of fragments assigned directly to this taxon
4. A rank code, indicating (U)nclassified, (R)oot, (D)omain, (K)ingdom,
   (P)hylum, (C)lass, (O)rder, (F)amily, (G)enus, or (S)pecies. Taxa that
   are not at any of these 10 ranks have a rank code that is formed by
   using the rank code of the closest ancestor rank with a number
   indicating the distance from that rank.  E.g., "G2" is a rank code
   indicating a taxon is between genus and species and the grandparent
   taxon is at the genus rank.
5. NCBI taxonomic ID number
6. Indented scientific name
```

###### Decontamination procedure

The decontamination procedure implemented in WGA-LP processes the sequencing reads directly and was built with the goal of eliminatig those reads that are confidently from the contaminant, avoiding loss of information. The following schema presents the decontamination algorithm:

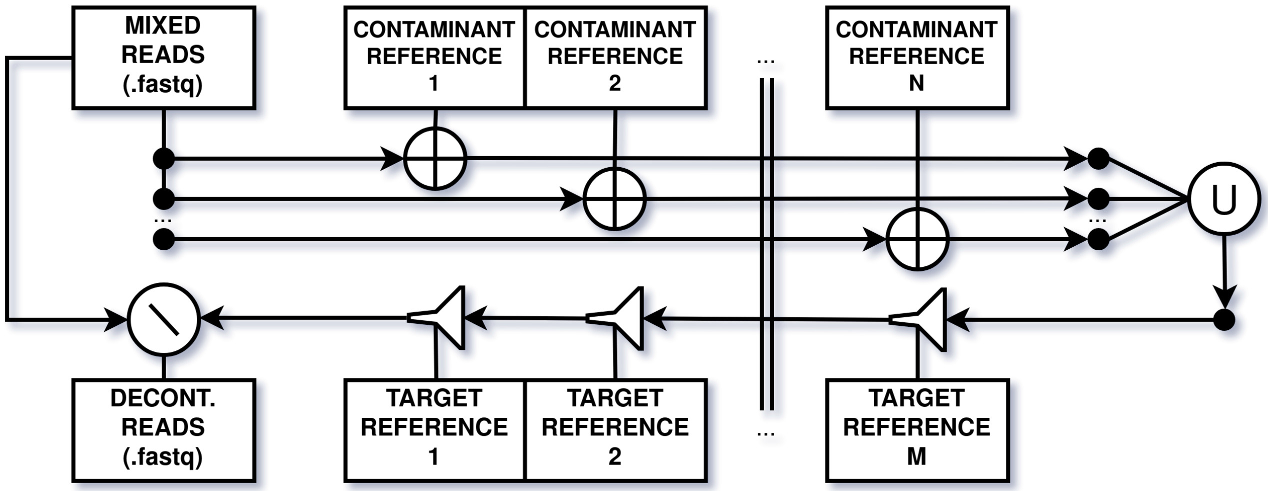

Input reads are mapped against each reference of the contaminant independently (first three wires from left to right). The mapped reads are then merged together (Union, ∪) and gradually filtered (last wire from right to left), with the effect of removing all the reads that map to any reference of the target organism. The final decontaminated reads are extracted by set difference (\) using the original input set. Each alignment step is composed by the subsequent application of three tools:

1. **bwa**: to align the reads to the references.
2. **samtools**: to filter and sort the alignment, in order to select mapped (or unmapped) reads.
3. **bazam**: to convert bam files of the alignment back into the fastq format.

In total, the procedure consists of a series of $N + M$ alignments, where $N$ is the number of the references for the target organism and $M$ the number of references of the contaminant. If needed, the algorithm can be run multiple times with different contaminants.

In order to run decontamination, we use the `wgalp decontaminate` command:

```
(base) wgalp:~/shared/144_working_directory>wgalp decontaminate
--- no arguments given, printing help message ---
Remove reads that map to a contaminant reference, but not to any target reference
references must be in .fasta format and BWA indexed (command: bwa index path/to/file.fasta)
--- arguments:
        --fastq : single raw read file to be cleaned (.fastq) [use only with NON paired end data]
        --fastq-fwd : forward raw reads (.fastq) [use only with paired end data]
        --fastq-rev : revese raw reads (.fastq) [use only with paired end data]
        --references : list of the possible references for the sequenced genome (.fasta, BWA indexed)
        --contaminants : list of the contaminant
        --output : the folder in which the output will be saved
        --help : print this message

(base) wgalp:~/shared/144_working_directory>
```

> Note that your references for the target and contaminant organisms **must** be indexed with bwa.
>
> That can be achieved with the command `bwa index file.fasta`
>
> If you have many references a `for` loop may be helpful:
>
> ```
> for f in `ls path/to/references/*.fasta`; do bwa index $f; done
> ```

In our example we run the decontamination as follows:

```
wgalp decontaminate \
    --fastq-fwd ../144/144_S13_L001_R1_001.fastq \
    --fastq-rev ../144/144_S13_L001_R2_001.fastq \
    --references ../references/rhamnosus/*.fasta \
    --contaminants ../references/pediococcus/*.fasta \
    --output decontamination
```

This step requires $M + N$ bwa alignments, where $M$ and $N$ are the number of references of the target organism and of the contaminant respectively :

```
[illegible]
force False
INFO: Step remove_good_reads_rev__affdf64f464580c8b00d2579902ad888
INFO: Running on {'fastq': 'decontamination/discarded_reads_rev.fastq', 'bam': 'decontamination/bad_reads__affdf64f464580c8b00d2579902ad888/aligned_to_Lrhamnosushsryfm1301.bam', '[illegible]
INFO:running >> samtools view decontamination/bad_reads__affdf64f464580c8b00d2579902ad888/aligned_to_Lrhamnosushsryfm1301.bam | cut -f 1 | uniq > decontamination/remove_good_reads_[illegible]
INFO: With outputs {'filtered_fastq': 'filtered_reads_remove_good_reads_rev__affdf64f464580c8b00d2579902ad888.fastq'}
task completed successfully
decontaminated reads are at the following locations:
        cleaned_fastq_fwd : decontamination/decontaminated_fwd.fastq
        cleaned_fastq_rev : decontamination/decontaminated_rev.fastq

real    49m9.513s
user    153m27.118s
sys     6m13.111s
wgalp:~/shared/144_working_directory>
```

We can check the resulting reads for contamination with `wgalp understand-origin`:

```
wgalp understand-origin \
    --fastq-fwd decontamination/decontaminated_fwd.fastq \
    --fastq-rev decontamination/decontaminated_rev.fastq \
    --kraken-db $kraken_db \
    --output kraken_after_decontamination
```

Kraken2 still show some reads from Pediococcus. Those reads still remains as Pediococcus and L. Rhamnosus are similar and there are valid mappings of them to the references of both target organism (L. Rhamnosus) and contaminant (Pediococcus).

```
...
71.33   580110   25051    G        1578                         Lactobacillus
...
0.66   5403     222      G        1253                         Pediococcus
...
```

Notice a drastic reduction of contaminant reads.

##### Run an assembler

WGA-LP supports SPAdes (plus SPAdes-Plasmid) and Minia assemblers natively and provides interfaces for their executions through the `wgalp assemble` command.

```
(base) wgalp:~/shared/144_working_directory>wgalp assemble
--- no arguments given, printing help message ---
Wrapper to easily run Whole Genome Assemblers
--- arguments:
        --fastq-fwd : preprocessed forward reads (.fastq)
        --fastq-rev : preprocessed reverse reads (.fastq)
        --output : path to the output folder
        --assembler : assembler name (from this list) [SPAdes, minia, SPAdes_plasmid]
        --kmer : kmer length (to be used only with minia)
        --help : print this message

(base) wgalp:~/shared/144_working_directory>
```

> Note that a user willing to use a **different assembler** can do so by:
>
> - running the assembler with its specific command on the decontaminated (or just trimmed) reads produced in the previous steps of WGA-LP
> - Using the nodes produced by the chosen assembler to the rest of the pipeline (scaffolds or contigs must be in **.fasta** format, comment lines will be used as unique IDs of nodes)

SPAdes tends to be a common choice for bacterial WGA, while Minia is a very simple and fast assembler that can be useful to evaluate the quality of the data.

In our example we use SPAdes to assemble the decontaminated reads:

```
wgalp assemble \
    --assembler SPAdes \
    --fastq-fwd decontamination/decontaminated_fwd.fastq \
    --fastq-rev decontamination/decontaminated_rev.fastq \
    --output SPAdes
```

with the following output:

```
[illegible]

Thank you for using SPAdes!
INFO: With outputs {'contigs': 'contigs.fasta', 'scaffolds': 'scaffolds.fasta', 'assembly_graph': 'assembly_graph.fastg'}
task completed successfully
assembled contigs/scaffolds are at the following locations:
        contigs : SPAdes/SPAdes/contigs.fasta
        scaffolds : SPAdes/SPAdes/scaffolds.fasta
check SPAdes for assembler specific files

real    12m43.011s
user    114m33.941s
sys     2m20.334s
(base) wgalp:~/shared/144_working_directory>
```

We can now start evaluating the quality of the nodes of the assembly.

##### Compute coverage statistics

An important step to check the quality of the final assembly is to realign reads to the assembly itself, in order to check the actual coverage of the produced nodes. `wgalp check-coverage` relies on bwa and `samtools depth` to create a summary of the coverages and length of each node:

```
(base) wgalp:~/shared/144_working_directory>wgalp check-coverage
--- no arguments given, printing help message ---
Program to extract coverage statistics from a Whole Genome Assembly
--- arguments:
        --fastq-fwd : raw forward reads (.fastq)
        --fastq-rev : raw reverse reads (.fastq)
        --contigs : assembled contigs (.fasta)
        --output : output folder
        --help : print this message

(base) wgalp:~/shared/144_working_directory>
```

In our example:

```
wgalp check-coverage \
    --fastq-fwd decontamination/decontaminated_fwd.fastq \
    --fastq-rev decontamination/decontaminated_rev.fastq \
    --contigs SPAdes/SPAdes/scaffolds.fasta \
    --output coverage
```

With the following output:

```
force False
INFO: Step samtools_depth
INFO: Running on {'input_bam': 'coverage/vsi_realign_contigs_depth/aligned_to_scaffolds.bam'}
INFO:running >> samtools depth coverage/vsi_realign_contigs_depth/aligned_to_scaffolds.bam  > coverage/samtools_depth/aligned_to_scaffolds.depth
INFO: With outputs {'depth_file': 'aligned_to_scaffolds.depth', 'depth_summary': 'aligned_to_scaffolds.depth.summary'}
task completed successfully
the file with the read depth for each base of the assembled genome is available at this location:
        depth_file : coverage/samtools_depth/aligned_to_scaffolds.depth
        depth_summary : coverage/samtools_depth/aligned_to_scaffolds.depth.summary

real    1m14.286s
user    5m59.634s
sys     0m15.579s
(base) wgalp:~/shared/144_working_directory>
```

The results will then be analyzed with WGA-LP as shown in the following sections.

##### Visualize coverage distribution

To check the quality of specific nodes by looking at the read pileup, it is possible to use `wgalp view-coverage` command as follows:

```
(base) wgalp:~/shared/144_working_directory>wgalp view-nodes
--- no arguments given, printing help message ---
Program to compute coverage plots for specific nodes from a samtools depth file
--- arguments:
        --depth : a .depth file from samtools depth
        --nodes : list of node IDs to be considered (node IDs are the same of the original fasta file of the assembly)
        --all : plot coverage for every node, overrides --nodes
        --output : output folder
        --help : print this message

(base) wgalp:~/shared/144_working_directory>
```

This is helpful to find anomalies in the coverage that may need further evaluation.

In our example:

```
wgalp view-nodes \
    --depth coverage/samtools_depth/aligned_to_scaffolds.depth \
    --all \
    --output coverage_plots
```

> The plotting can be limited with specific IDs with **–nodes** flag

with output:

```
Saving 7 x 7 in image
[1] "preparing graph for NODE_1628_length_133_cov_1.333333 ..."
Saving 7 x 7 in image
[1] "preparing graph for NODE_1629_length_128_cov_2633.000000 ..."
Saving 7 x 7 in image
INFO: With outputs {'output_dir': ''}
task completed successfully
the plots of with coverages were saved at this location:
        output_dir : coverage_plots/coverage_plots/

real    14m51.714s
user    14m40.687s
sys     0m12.788s
(base) wgalp:~/shared/144_working_directory>
```

This analysis can be postponed also after a first node selection, to widely reduce the produced plots.

This is an example of an output plot from the current test:

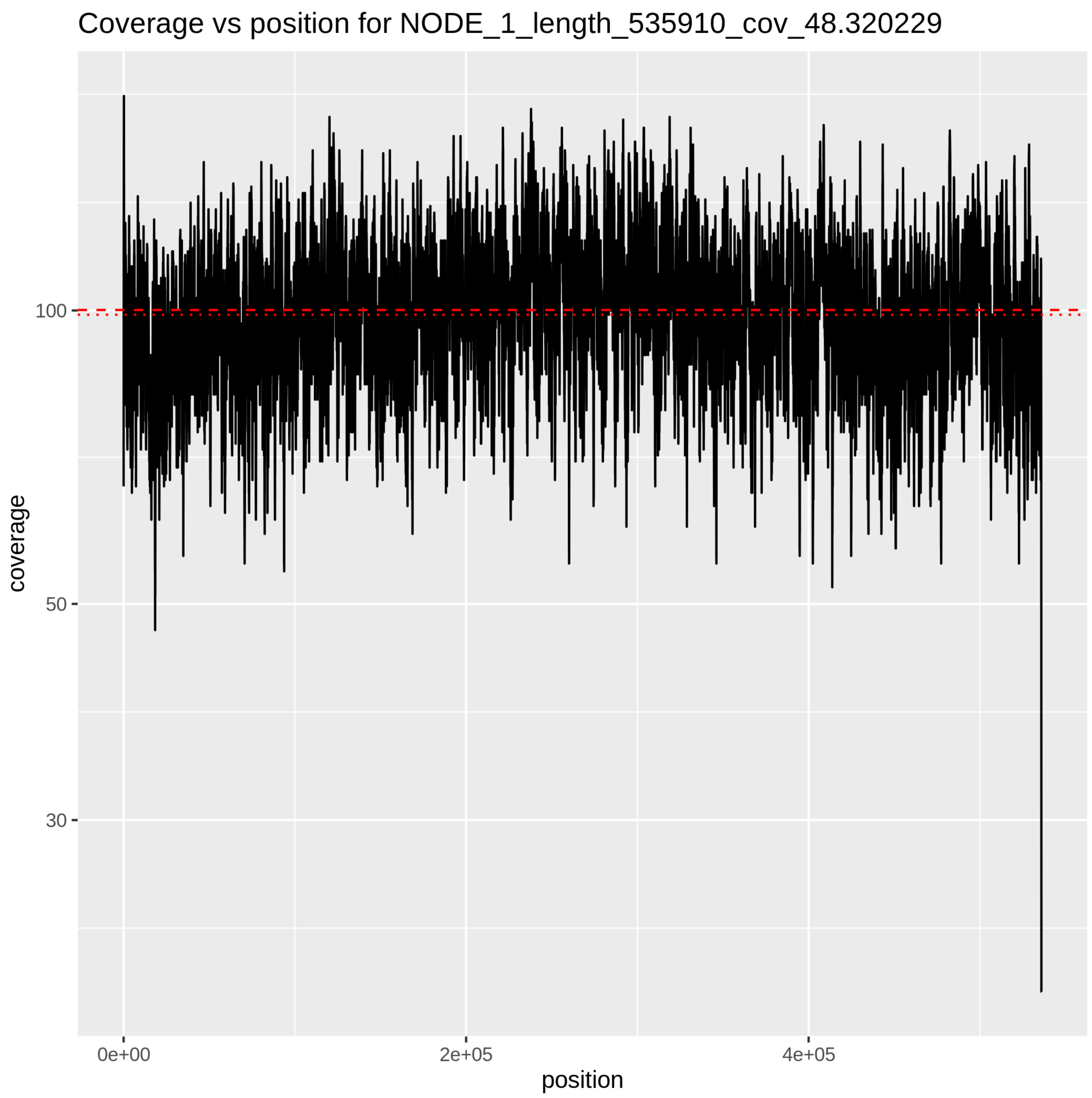

A more interesting example is the 6th node of the assembly:

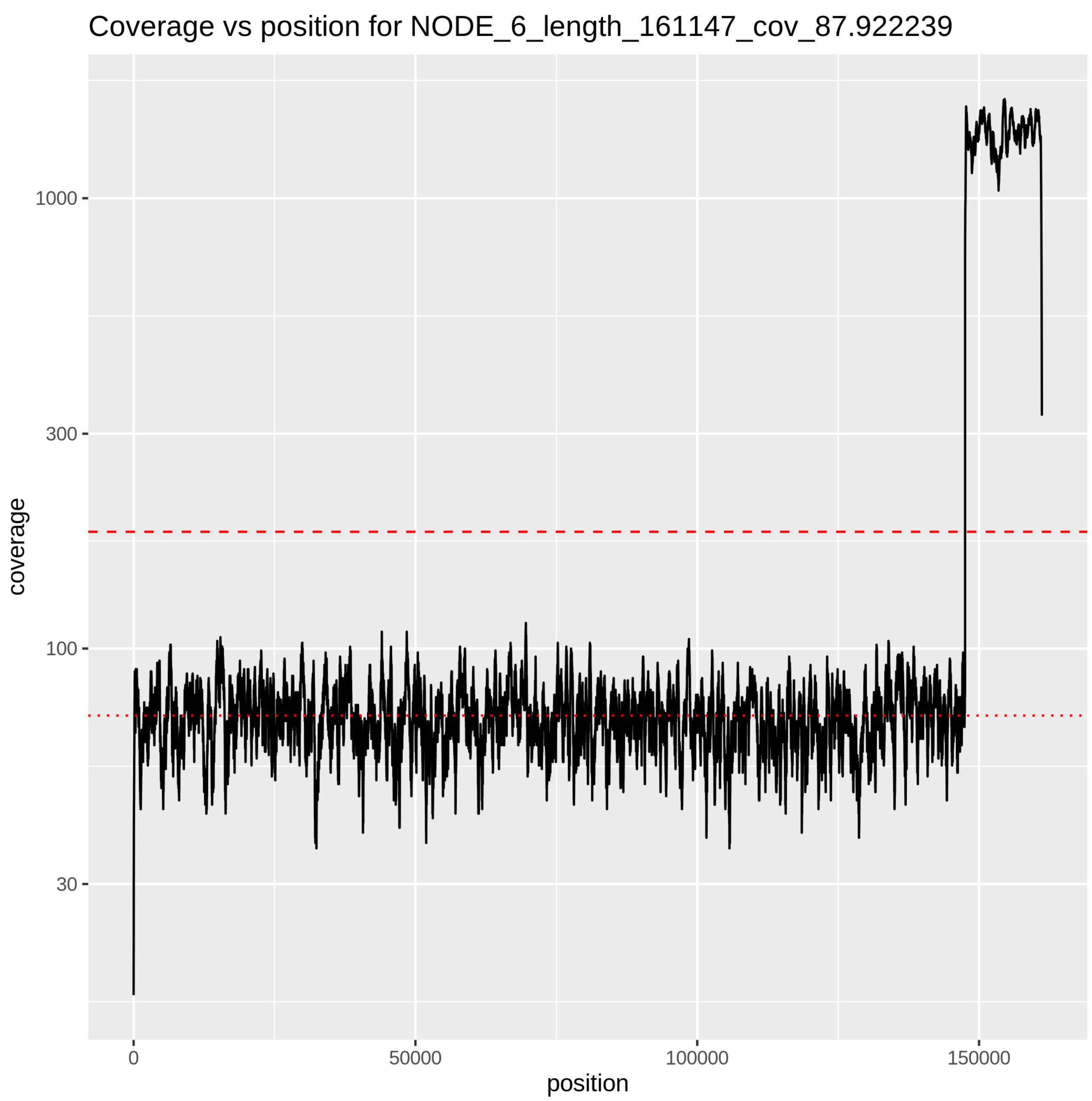

That includes a peak of coverage at its end. Using `blast` on the higher coverage portion we get the following report:

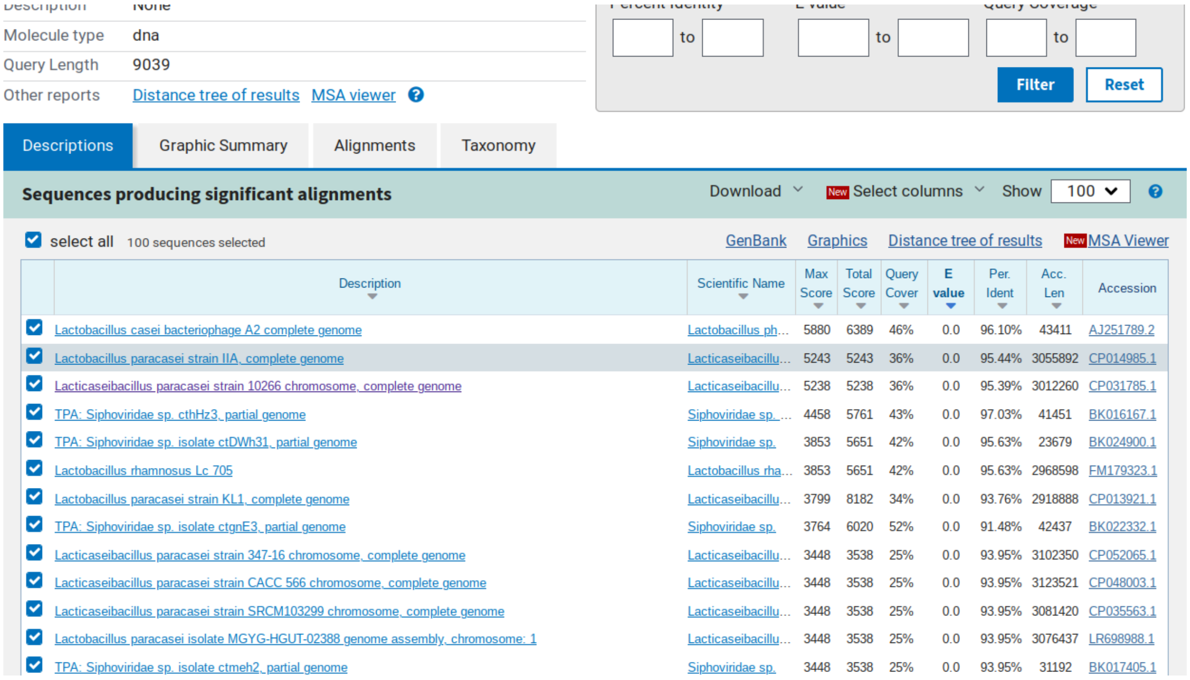

suggesting that a possible cause of the coverage peak is the insertion of a bacteriophage genome in the bacterial genome.

##### Visualize nodes by length and coverage

To have a better understanding of the characteristics of the nodes produced by the assembler, we developed a web app that is capable of visualizing the `.depth.summmary` output of `wgalp check-coverage`. The web app is very simple and needs just the upload of the `.depth.summary` file.

Link: https://redsnic.shinyapps.io/ContigCoverageVisualizer/

Mantainer: Nicolò Rossi

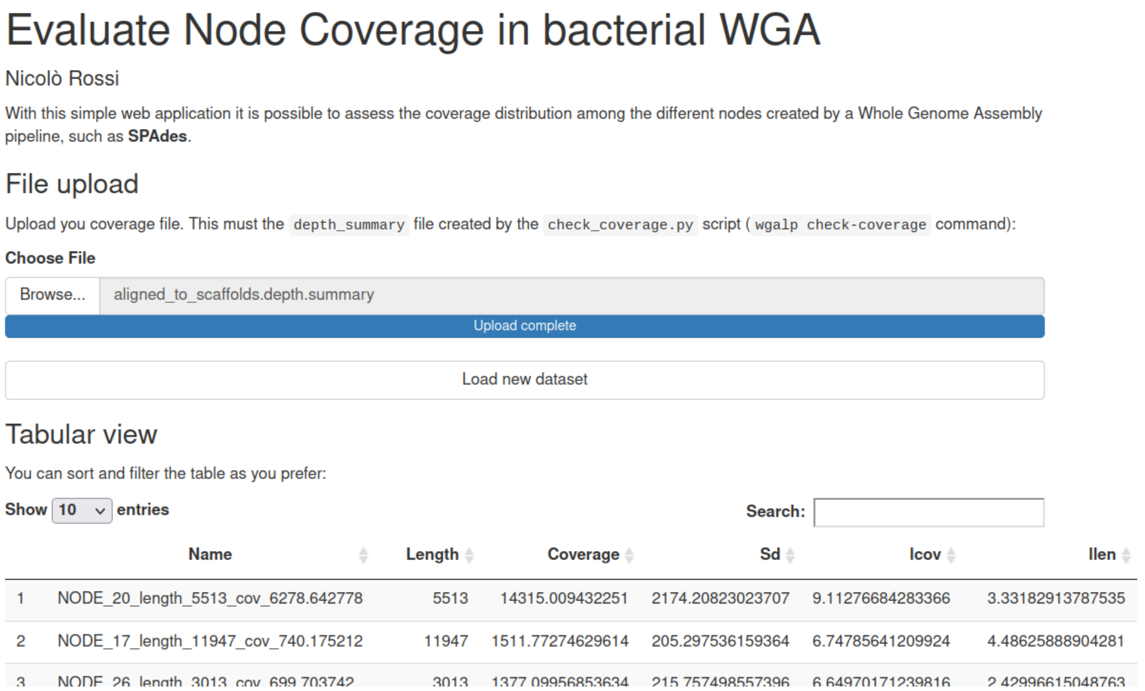

> **No information** is saved about the analysis done with this web app.

The distribution of nodes in term of length and coverage is visualized in an interactive scatterplot. Hovering on nodes with the mouse shows the name of the node and its position as in this image (in the example, target coverage was 100x):

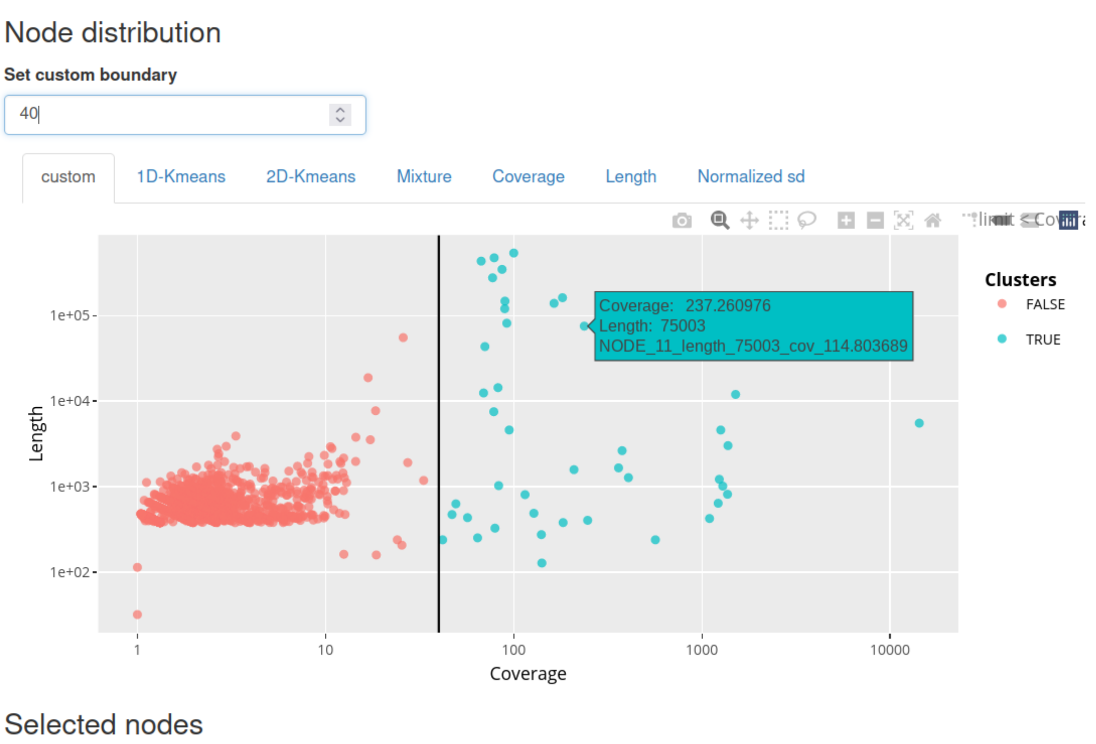

The selection of nodes is always based on the simple "Custom" boundary over coverages. The other tabs show useful clustering methods and distributions. Selection always keeps the nodes over the boundary.

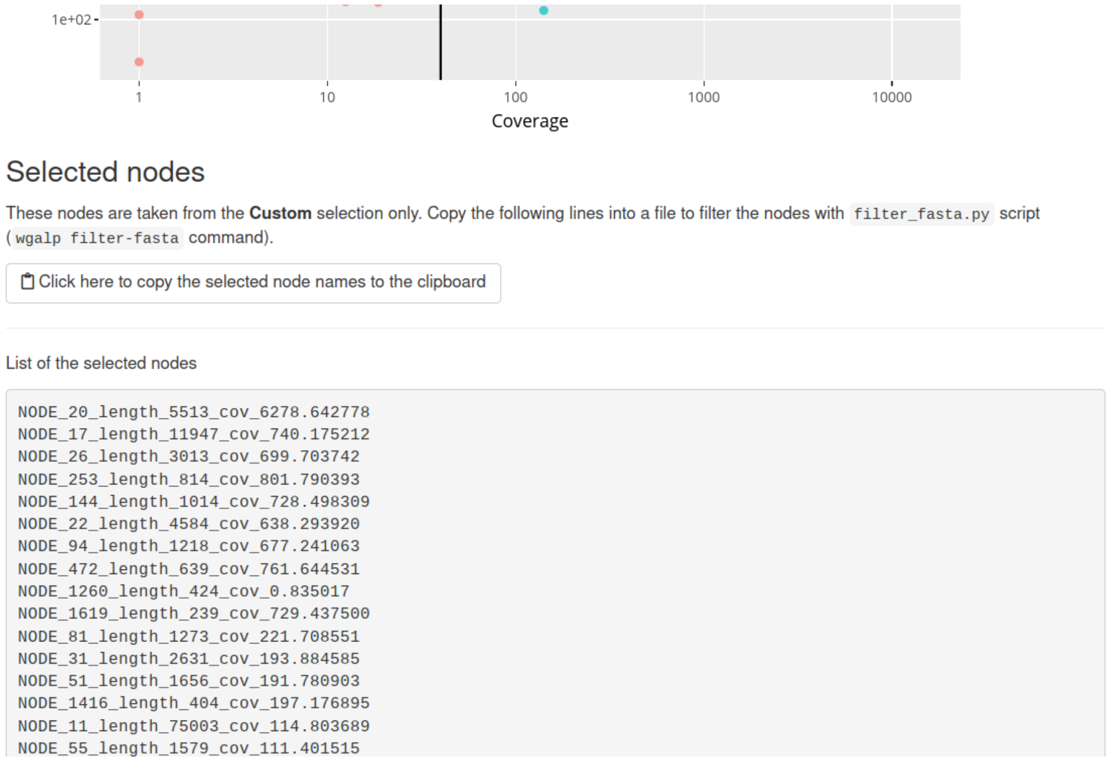

Using the "copy to clipboard" button, it is possible to get the selected node IDs for further use with `wgalp`

`filter-assembly`.

Finally, it is possible to evaluate the effects of the selection on clusterization and distribution. This can help in finding the next nodes to investigate:

```
NODE_1181_length_434_cov_0.768730
NODE_491_length_629_cov_39.705179
NODE_885_length_472_cov_2.556522
NODE_1620_length_239_cov_37.741071
```

After selection

Check the status after the application of the custom boundary. This can be useful to refine the analysis.

1D-Kmeans | 2D-Kmeans | Mixture | Coverage | Length | Normalized sd

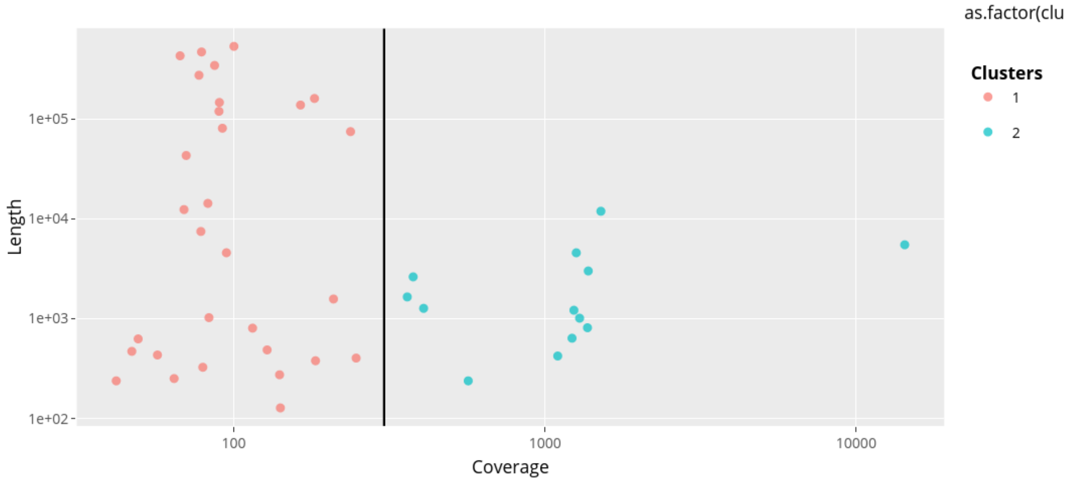

###### Filter nodes from the assembly

We can use "copy and paste" with the `cat` command (or any file editor, even in the host system if using `Docker`) to create a file whit the IDs of the selected nodes:

```
(base) wgalp:~/shared/144_working_directory>cat > selection40.txt
NODE_20_length_5513_cov_6278.642778
NODE_17_length_11947_cov_740.175212
NODE_26_length_3013_cov_699.703742
NODE_253_length_814_cov_801.790393
NODE_144_length_1014_cov_728.498309
NODE_22_length_4584_cov_638.293920
NODE_94_length_1218_cov_677.241063
NODE_472_length_639_cov_761.644531
NODE_1260_length_424_cov_0.835017
NODE_1619_length_239_cov_729.437500
NODE_81_length_1273_cov_221.708551
NODE_31_length_2631_cov_193.884585
NODE_51_length_1656_cov_191.780903
NODE_1416_length_404_cov_197.176895
NODE_11_length_75003_cov_114.803689
NODE_55_length_1579_cov_111.401515
NODE_1604_length_380_cov_0.920949
NODE_6_length_161147_cov_87.922239
NODE_8_length_138419_cov_80.429497
NODE_1629_length_128_cov_2633.000000
NODE_1617_length_275_cov_143.182432
NODE_828_length_488_cov_86.554017
NODE_261_length_806_cov_66.578792
NODE_1_length_535910_cov_48.320229
NODE_21_length_4587_cov_90.636547
NODE_10_length_81116_cov_44.490511
NODE_7_length_146963_cov_43.291046
NODE_9_length_119768_cov_43.192835
NODE_4_length_345213_cov_41.849811
NODE_140_length_1027_cov_47.310000
NODE_15_length_14331_cov_40.005210
NODE_1616_length_327_cov_41.765000
NODE_2_length_471473_cov_37.993595
NODE_19_length_7498_cov_35.392077
NODE_5_length_275244_cov_37.938339
NODE_13_length_43256_cov_34.005055
NODE_16_length_12423_cov_33.704863
NODE_3_length_431246_cov_32.404519
NODE_1618_length_252_cov_85.104000
NODE_1181_length_434_cov_0.768730
NODE_491_length_629_cov_39.705179
NODE_885_length_472_cov_2.556522
NODE_1620_length_239_cov_37.741071
(base) wgalp:~/shared/144_working_directory>
```

Then, it is possible to proceed with the actual selection:

```
(base) wgalp:~/shared/144_working_directory>wgalp filter-assembly
--- no arguments given, printing help message ---
Select nodes by ID from a Whole Genom Assembly
--- arguments:
        --contigs : assembled contigs or scaffolds (.fasta)
        --selected-contigs : a file containing the ids of the selected contigs (each id is in its own line)
        --complement : if set, keeps contigs not in list
        --output : path to the output folder
        --help : print this message

(base) wgalp:~/shared/144_working_directory>
```

In our example:

```
wgalp filter-assembly \
    --contigs SPAdes/SPAdes/scaffolds.fasta \
    --selected-contigs selection40.txt \
    --output filtered_contigs
```

```
(base) wgalp:~/shared/144_working_directory>time wgalp filter-assembly --contigs SPAdes/SPAdes/scaffolds.fasta --selected-contigs selection40.txt --output filtered_contigs
task completed successfully
filtered .fasta is at the followinf location:
        filtered_fasta : filtered_contigs/filtered_contigs.fasta

real    0m0.783s
user    0m1.095s
sys     0m1.255s
(base) wgalp:~/shared/144_working_directory>
```

> **–complement** flag allows the user to select for the discarded nodes. Using this option may allow to check that no possibly valid node is discarded.

###### Refine node selection

Now it is **highly recommended** to use Kraken2 (through `wgalp understand-origin`) to check if there are nodes that are assemblies of reads of the contaminant. Such prediction must be evaluated also with **blast**.

In our example:

```
wgalp understand-origin \
    --fasta filtered_contigs/filtered_contigs.fasta \
    --kraken-db $kraken_db \
    --output node_origin
```

using the command:

```
cat node_origin/kraken/kraken.log | cut -d$'\t' -f 2,3
```

we get the following table (note that coverages in the names are computed by SPAdes using a different approach then read-remapping):

```
NODE_1_length_535910_cov_48.320229    Lactobacillus rhamnosus ...
NODE_2_length_471473_cov_37.993595    Lactobacillus rhamnosus ...
NODE_3_length_431246_cov_32.404519    Lactobacillus rhamnosus ...
NODE_4_length_345213_cov_41.849811    Lactobacillus rhamnosus ...
NODE_5_length_275244_cov_37.938339    Lactobacillus rhamnosus ...
```

```
NODE_6_length_161147_cov_87.922239    Lactobacillus rhamnosus ...
NODE_7_length_146963_cov_43.291046    Lactobacillus rhamnosus ...
NODE_8_length_138419_cov_80.429497    Lactobacillus rhamnosus ...
NODE_9_length_119768_cov_43.192835    Lactobacillus rhamnosus ...
NODE_10_length_81116_cov_44.490511    Lactobacillus rhamnosus ...
NODE_11_length_75003_cov_114.803689   Lactobacillus paracasei ...
NODE_13_length_43256_cov_34.005055    Lactobacillus rhamnosus ...
NODE_15_length_14331_cov_40.005210    Lactobacillus rhamnosus ...
NODE_16_length_12423_cov_33.704863    Lactobacillus rhamnosus ...
NODE_17_length_11947_cov_740.175212   Lactobacillus rhamnosus ...
NODE_19_length_7498_cov_35.392077     Pediococcus acidilactici ...
NODE_20_length_5513_cov_6278.642778   Bacteria ...
NODE_21_length_4587_cov_90.636547     Lactobacillus rhamnosus ...
NODE_22_length_4584_cov_638.293920    Lactobacillus rhamnosus ...
NODE_26_length_3013_cov_699.703742    Lactobacillus paracasei ...
NODE_31_length_2631_cov_193.884585    Lactobacillus rhamnosus ...
NODE_51_length_1656_cov_191.780903    Lactobacillus rhamnosus ...
NODE_55_length_1579_cov_111.401515    Lactobacillus rhamnosus ...
NODE_81_length_1273_cov_221.708551    Lactobacillus rhamnosus ...
NODE_94_length_1218_cov_677.241063    Lactobacillus rhamnosus ...
NODE_140_length_1027_cov_47.310000    Lactobacillus rhamnosus ...
NODE_144_length_1014_cov_728.498309   Lactobacillus rhamnosus ...
NODE_253_length_814_cov_801.790393    Lactobacillus rhamnosus ...
NODE_261_length_806_cov_66.578792     Lactobacillus casei group ...
NODE_472_length_639_cov_761.644531    Lactobacillus rhamnosus ...
NODE_491_length_629_cov_39.705179     Lactobacillus rhamnosus ...
NODE_828_length_488_cov_86.554017     Lactobacillus rhamnosus ...
NODE_885_length_472_cov_2.556522      Pediococcus acidilactici ...
NODE_1181_length_434_cov_0.768730     Lactobacillus rhamnosus ...
NODE_1260_length_424_cov_0.835017     cellular organisms ...
NODE_1416_length_404_cov_197.176895   Lactobacillus ...
NODE_1604_length_380_cov_0.920949     Bacteria ...
NODE_1616_length_327_cov_41.765000    Lactobacillus rhamnosus ...
NODE_1617_length_275_cov_143.182432   Lactobacillus rhamnosus ...
NODE_1618_length_252_cov_85.104000    Lactobacillus rhamnosus ...
NODE_1619_length_239_cov_729.437500   Lactobacillus paracasei ...
NODE_1620_length_239_cov_37.741071    Lactobacillales ...
NODE_1629_length_128_cov_2633.000000  unclassified ...
```

We see that there are many short nodes, moreover some scaffolds are labeled as derived from Pediococcus, the contaminant. Shorter reads may be evaluated as they can be valid Insertion Sequences (IS) that are hard to assemble. In this example we focus on three nodes:

```
NODE_19_length_7498_cov_35.392077      Pediococcus acidilactici (taxid 1254)
NODE_20_length_5513_cov_6278.642778    Bacteria (taxid 2)
```

```
NODE_885_length_472_cov_2.556522        Pediococcus acidilactici (taxid 1254)
```

And we use **blast** to see if kraken2 prediction are or not reliable. To get the sequences, we can either search the `.fasta` file containing the nodes directly with a file editor, or use `wgalp filter-assembly` command.

`NODE_20_length_5513_cov_6278.642778` is the $\Phi X147$:

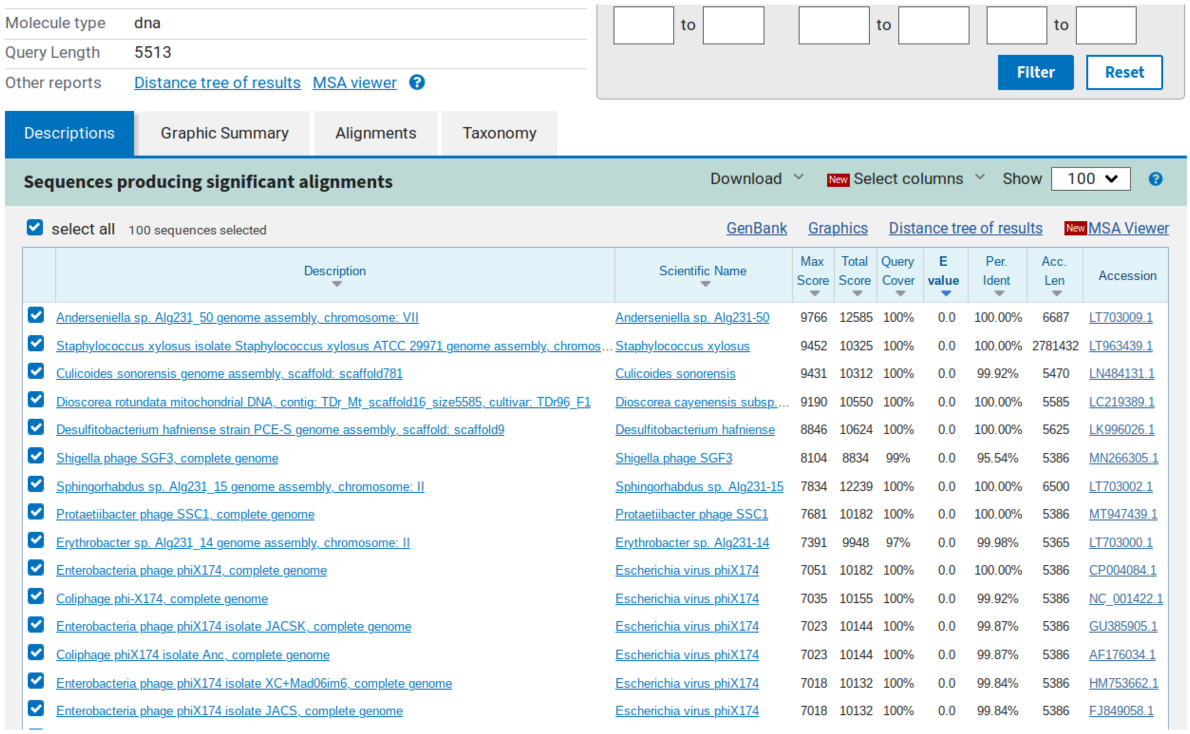

Molecule type: dna
Query Length: 5513
Other reports: Distance tree of results MSA viewer

to | to | to | Filter | Reset

Descriptions | Graphic Summary | Alignments | Taxonomy

**Sequences producing significant alignments** Download Select columns Show 100

select all 100 sequences selected GenBank Graphics Distance tree of results MSA Viewer

| Description | Scientific Name | Max Score | Total Score | Query Cover | E value | Per. Ident | Acc. Len | Accession |
|---|---|---|---|---|---|---|---|---|
| Anderseniella sp. Alg231_50 genome assembly, chromosome: VII | Anderseniella sp. Alg231-50 | 9766 | 12585 | 100% | 0.0 | 100.00% | 6687 | LT703009.1 |
| Staphylococcus xylosus isolate Staphylococcus xylosus ATCC 29971 genome assembly, chromos... | Staphylococcus xylosus | 9452 | 10325 | 100% | 0.0 | 100.00% | 2781432 | LT963439.1 |
| Culicoides sonorensis genome assembly, scaffold: scaffold781 | Culicoides sonorensis | 9431 | 10312 | 100% | 0.0 | 99.92% | 5470 | LN484131.1 |
| Dioscorea rotundata mitochondrial DNA, contig: TDr_Mt_scaffold16_size5585, cultivar: TDr96_F1 | Dioscorea cayenensis subsp.... | 9190 | 10550 | 100% | 0.0 | 100.00% | 5585 | LC219389.1 |
| Desulfitobacterium hafniense strain PCE-S genome assembly, scaffold: scaffold9 | Desulfitobacterium hafniense | 8846 | 10624 | 100% | 0.0 | 100.00% | 5625 | LK996026.1 |
| Shigella phage SGF3, complete genome | Shigella phage SGF3 | 8104 | 8834 | 99% | 0.0 | 95.54% | 5386 | MN266305.1 |
| Sphingorhabdus sp. Alg231_15 genome assembly, chromosome: II | Sphingorhabdus sp. Alg231-15 | 7834 | 12239 | 100% | 0.0 | 100.00% | 6500 | LT703002.1 |
| Protaetiibacter phage SSC1, complete genome | Protaetiibacter phage SSC1 | 7681 | 10182 | 100% | 0.0 | 100.00% | 5386 | MT947439.1 |
| Erythrobacter sp. Alg231_14 genome assembly, chromosome: II | Erythrobacter sp. Alg231-14 | 7391 | 9948 | 97% | 0.0 | 99.98% | 5365 | LT703000.1 |
| Enterobacteria phage phiX174, complete genome | Escherichia virus phiX174 | 7051 | 10182 | 100% | 0.0 | 100.00% | 5386 | CP004084.1 |
| Coliphage phi-X174, complete genome | Escherichia virus phiX174 | 7035 | 10155 | 100% | 0.0 | 99.92% | 5386 | NC_001422.1 |
| Enterobacteria phage phiX174 isolate JACSK, complete genome | Escherichia virus phiX174 | 7023 | 10144 | 100% | 0.0 | 99.87% | 5386 | GU385905.1 |
| Coliphage phiX174 isolate Anc, complete genome | Escherichia virus phiX174 | 7023 | 10144 | 100% | 0.0 | 99.87% | 5386 | AF176034.1 |
| Enterobacteria phage phiX174 isolate XC+Mad06im6, complete genome | Escherichia virus phiX174 | 7018 | 10132 | 100% | 0.0 | 99.84% | 5386 | HM753662.1 |
| Enterobacteria phage phiX174 isolate JACS, complete genome | Escherichia virus phiX174 | 7018 | 10132 | 100% | 0.0 | 99.84% | 5386 | FJ849058.1 |

a phage used in Illumina sequencing as reference. This node must be removed.

`NODE_19_length_7498_cov_35.392077` seems to actually be a fragment of the contaminant's genome:

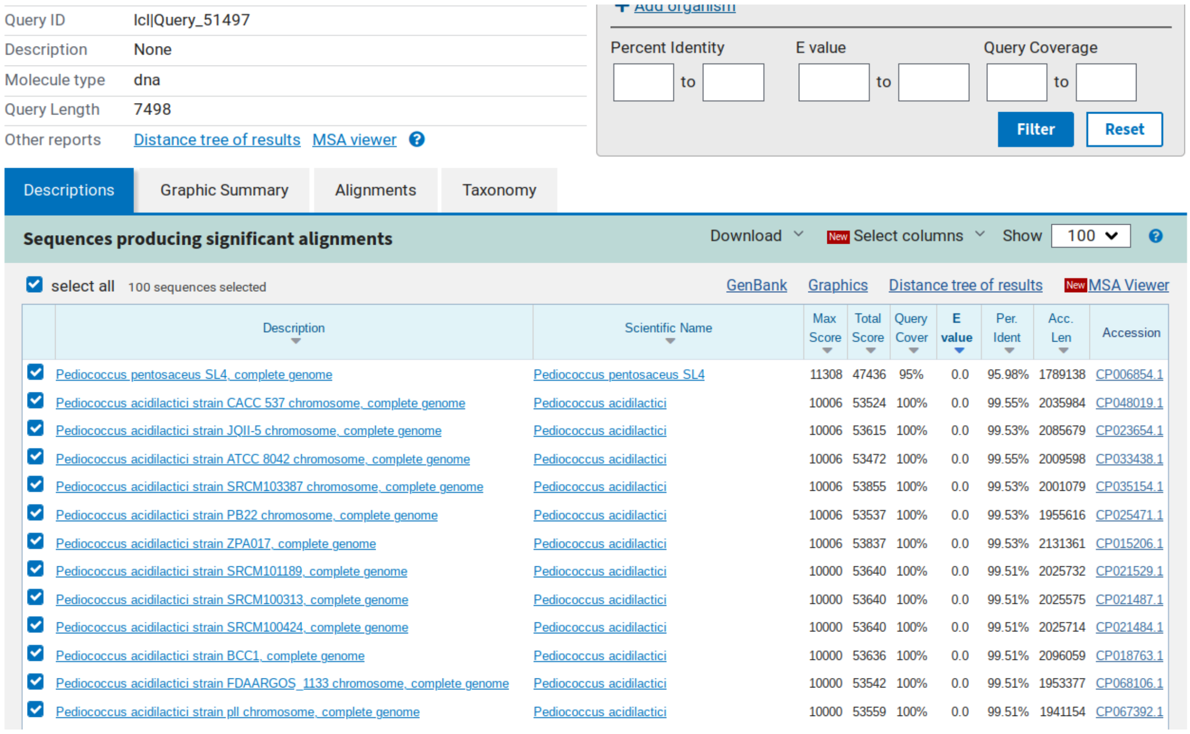

Query ID: lcl|Query_51497
Description: None
Molecule type: dna
Query Length: 7498
Other reports: Distance tree of results MSA viewer

Add organism

Percent Identity: to | E value: to | Query Coverage: to | Filter | Reset

Descriptions | Graphic Summary | Alignments | Taxonomy

**Sequences producing significant alignments** Download Select columns Show 100

select all 100 sequences selected GenBank Graphics Distance tree of results MSA Viewer

| Description | Scientific Name | Max Score | Total Score | Query Cover | E value | Per. Ident | Acc. Len | Accession |
|---|---|---|---|---|---|---|---|---|
| Pediococcus pentosaceus SL4, complete genome | Pediococcus pentosaceus SL4 | 11308 | 47436 | 95% | 0.0 | 95.98% | 1789138 | CP006854.1 |
| Pediococcus acidilactici strain CACC 537 chromosome, complete genome | Pediococcus acidilactici | 10006 | 53524 | 100% | 0.0 | 99.55% | 2035984 | CP048019.1 |
| Pediococcus acidilactici strain JQII-5 chromosome, complete genome | Pediococcus acidilactici | 10006 | 53615 | 100% | 0.0 | 99.53% | 2085679 | CP023654.1 |
| Pediococcus acidilactici strain ATCC 8042 chromosome, complete genome | Pediococcus acidilactici | 10006 | 53472 | 100% | 0.0 | 99.55% | 2009598 | CP033438.1 |
| Pediococcus acidilactici strain SRCM103387 chromosome, complete genome | Pediococcus acidilactici | 10006 | 53855 | 100% | 0.0 | 99.53% | 2001079 | CP035154.1 |
| Pediococcus acidilactici strain PB22 chromosome, complete genome | Pediococcus acidilactici | 10006 | 53537 | 100% | 0.0 | 99.53% | 1955616 | CP025471.1 |
| Pediococcus acidilactici strain ZPA017, complete genome | Pediococcus acidilactici | 10006 | 53837 | 100% | 0.0 | 99.53% | 2131361 | CP015206.1 |
| Pediococcus acidilactici strain SRCM101189, complete genome | Pediococcus acidilactici | 10000 | 53640 | 100% | 0.0 | 99.51% | 2025732 | CP021529.1 |
| Pediococcus acidilactici strain SRCM100313, complete genome | Pediococcus acidilactici | 10000 | 53640 | 100% | 0.0 | 99.51% | 2025575 | CP021487.1 |
| Pediococcus acidilactici strain SRCM100424, complete genome | Pediococcus acidilactici | 10000 | 53640 | 100% | 0.0 | 99.51% | 2025714 | CP021484.1 |
| Pediococcus acidilactici strain BCC1, complete genome | Pediococcus acidilactici | 10000 | 53636 | 100% | 0.0 | 99.51% | 2096059 | CP018763.1 |
| Pediococcus acidilactici strain FDAARGOS_1133 chromosome, complete genome | Pediococcus acidilactici | 10000 | 53542 | 100% | 0.0 | 99.51% | 1953377 | CP068106.1 |
| Pediococcus acidilactici strain pll chromosome, complete genome | Pediococcus acidilactici | 10000 | 53559 | 100% | 0.0 | 99.51% | 1941154 | CP067392.1 |

It is likely that this node is actually a small assembly of the contaminant. We will remove it.

Similarly, also `NODE_885_length_472_cov_2.556522` is confirmed as originally from the contaminant by blast.

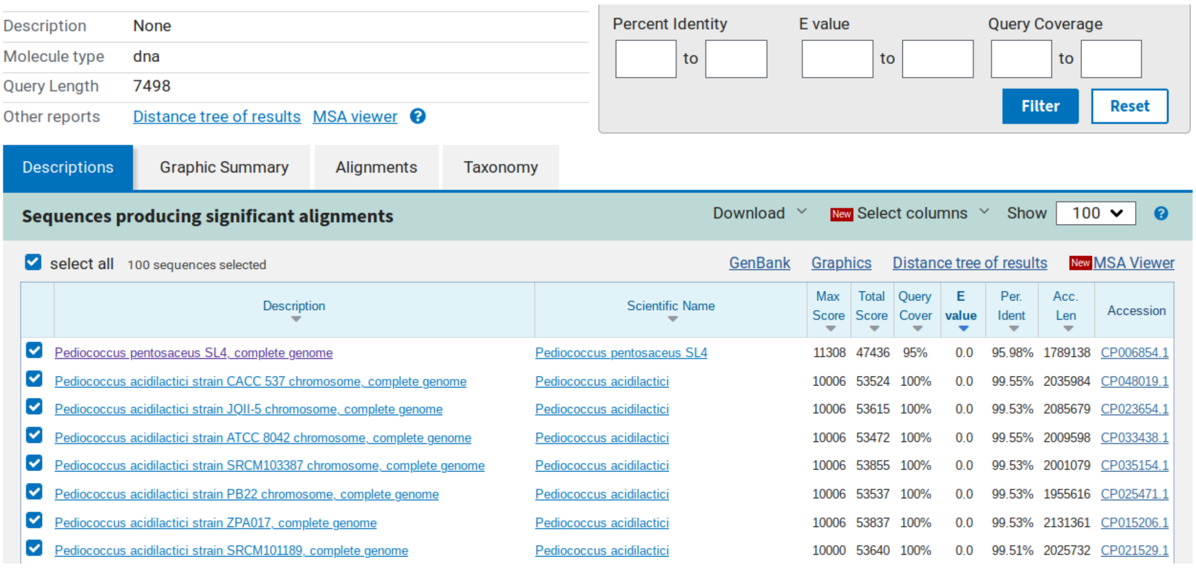

| | |
|---|---|
| Description | None |
| Molecule type | dna |
| Query Length | 7498 |
| Other reports | Distance tree of results MSA viewer |

Percent Identity to | E value to | Query Coverage to | Filter | Reset

Descriptions | Graphic Summary | Alignments | Taxonomy

**Sequences producing significant alignments** Download | New Select columns | Show 100

select all 100 sequences selected | GenBank | Graphics | Distance tree of results | New MSA Viewer

| Description | Scientific Name | Max Score | Total Score | Query Cover | E value | Per. Ident | Acc. Len | Accession |
|---|---|---|---|---|---|---|---|---|
| Pediococcus pentosaceus SL4, complete genome | Pediococcus pentosaceus SL4 | 11308 | 47436 | 95% | 0.0 | 95.98% | 1789138 | CP006854.1 |
| Pediococcus acidilactici strain CACC 537 chromosome, complete genome | Pediococcus acidilactici | 10006 | 53524 | 100% | 0.0 | 99.55% | 2035984 | CP048019.1 |
| Pediococcus acidilactici strain JQII-5 chromosome, complete genome | Pediococcus acidilactici | 10006 | 53615 | 100% | 0.0 | 99.53% | 2085679 | CP023654.1 |
| Pediococcus acidilactici strain ATCC 8042 chromosome, complete genome | Pediococcus acidilactici | 10006 | 53472 | 100% | 0.0 | 99.55% | 2009598 | CP033438.1 |
| Pediococcus acidilactici strain SRCM103387 chromosome, complete genome | Pediococcus acidilactici | 10006 | 53855 | 100% | 0.0 | 99.53% | 2001079 | CP035154.1 |
| Pediococcus acidilactici strain PB22 chromosome, complete genome | Pediococcus acidilactici | 10006 | 53537 | 100% | 0.0 | 99.53% | 1955616 | CP025471.1 |
| Pediococcus acidilactici strain ZPA017, complete genome | Pediococcus acidilactici | 10006 | 53837 | 100% | 0.0 | 99.53% | 2131361 | CP015206.1 |
| Pediococcus acidilactici strain SRCM101189, complete genome | Pediococcus acidilactici | 10000 | 53640 | 100% | 0.0 | 99.51% | 2025732 | CP021529.1 |

We can remove the unwanted nodes simply by editing the `.fasta` file, or by using `wgalp filter-assembly --complement`.

```
wgalp filter-assembly \
    --complement \
    --contigs filtered_contigs/filtered_contigs.fasta \
    --selected-contigs remove.nodes \
    --output precise_filter
```

> If needed, other node based filters should be applied at this step

##### Reorder the assembly using a reference genome

Using **Mauve** aligner, it is possible to optimally reorder nodes to follow a reference genome:

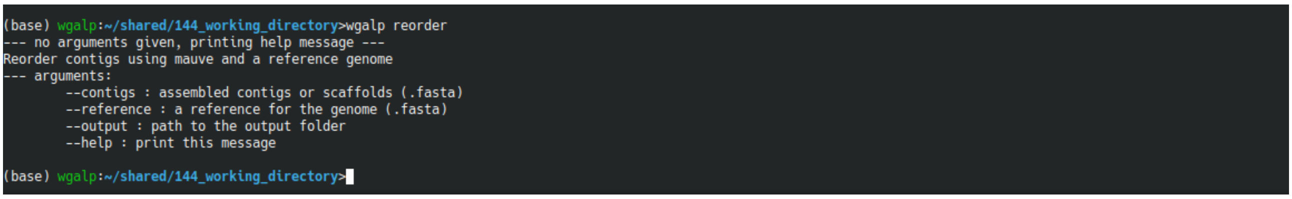

```
(base) wgalp:~/shared/144_working_directory>wgalp reorder
--- no arguments given, printing help message ---
Reorder contigs using mauve and a reference genome
--- arguments:
        --contigs : assembled contigs or scaffolds (.fasta)
        --reference : a reference for the genome (.fasta)
        --output : path to the output folder
        --help : print this message

(base) wgalp:~/shared/144_working_directory>
```

In our example:

```
wgalp reorder \
    --contigs precise_filter/filtered_contigs.fasta \
    --reference ../references/rhamnosus/LrhamnosusGGATCC.fasta \
    --output reordering
```

With the following output:

```
task completed successfully
the reordered assembly is at the following location:
        contigs : reordering/mauve_reorder/alignment2/filtered_contigs.fasta
other formats are available in the output folder reordering

real    1m0.135s
user    1m7.050s
sys     0m4.609s
wgalp:~/shared/144_working_directory>
```

##### Extract plasmids using Recycler

WGA-LP also includes two tools to extract putative plasmid, the first is **SPAdes plasmid** included in `wgalp assemble` command, the latter is **Recycler**. Recycler can be run with `wgalp plasmid` command:

```
(base) wgalp:~/shared/144_working_directory>wgalp plasmid
--- no arguments given, printing help message ---
Wrapper to easily run recycler and inferr plasmids from genome assembly graphs
--- arguments:
        --fastq-fwd : raw forward reads (.fastq)
        --fastq-rev : raw reverse reads (.fastq)
        --contigs : assembled contigs (.fasta)
        --assembly-graph : assembly graph (.fastg)
        --kmer : maximum kmer length used by the assembler (127 for spades)
        --output : output folder
        --help : print this message

(base) wgalp:~/shared/144_working_directory>
```

In our example:

```
wgalp plasmid \
    --fastq-fwd decontamination/decontaminated_fwd.fastq \
    --fastq-rev decontamination/decontaminated_rev.fastq \
    --contigs precise_filter/filtered_contigs.fasta \
    --assembly-graph SPAdes/SPAdes/assembly_graph.fastg \
    --kmer 127 \
    --output recycler
```

```
[illegible]
(67.333333, 656.1080362605982, 746.3048453999997)
================== path, coverage levels when added ====================
(200, ' nodes remain in component\n')
("EDGE_386451_length_171_cov_16.090909'", "EDGE_100090_length_16216_cov_12.190068'")
('before', [16.090909, 12.190068])
('after', [3.8730411870123476, 0])
(2, ' nodes remain in component\n')
==================final_paths identities after updates: ================
("EDGE_386451_length_171_cov_16.090909'", "EDGE_100090_length_16216_cov_12.190068'")

("EDGE_100070_length_5513_cov_6278.642778'",)

INFO: With outputs {'plasmid_fasta': 'assembly_graph.cycs.fasta'}
task completed successfully
the fasta file with the inferred plasmids is at the following location:
        plasmid_fasta : recycler_clean/recycler/assembly_graph.cycs.fasta

real    2m57.830s
user    6m6.910s
sys     0m18.906s
(base) wgalp:~/shared/144_working_directory>
```

This run leads to two putative plasmids.

```
>RNODE_2_length_16133_cov_12.42334
>RNODE_1_length_5386_cov_6278.64278
```

The first is a plasmid from the contaminant, while the second is genome of the $\Phi X174$ phage. There is no trace of these sequences in the final assembly (as they are cut when cleaning the nodes of the assembly). So in this case there are no putative plasmids for this genome.

> This can be easily checked using `blast` with the nodes in `assembly_graph.cycs.fasta`. To see that the sequences are absent in the assembled genome, it is possible to use `two sequence blast`.

##### Assess the quality of the final assembly

WGA-LP includes a suite of programs for the quality test of the resulting assembly. This includes **Quast**, **checkM**, and **Merqury**:

```
(base) wgalp:~/shared/144_working_directory>wgalp quality
--- no arguments given, printing help message ---
Run tools to evaluate WGA quality
--- arguments:
        --fastq-fwd : raw forward reads (.fastq)
        --fastq-rev : raw reverse reads (.fastq)
        --assembly : WGA assembly to evaluate (.fasta)
        --output : path to the output folder
        --full-tree : use full tree in checkM instead reduced_tree (requires > 40GB of ram)
        --kmer-length : kmer size to be used in merqury (use 16 for 3Mpb, check with: $MERQURY/best_k.sh <genome_size>)
        --help : print this message

(base) wgalp:~/shared/144_working_directory>
```

In our example:

```
wgalp quality \
    --fastq-fwd decontamination/decontaminated_fwd.fastq \
    --fastq-rev decontamination/decontaminated_rev.fastq \
    --assembly reordering/mauve_reorder/alignment2/filtered_contigs.fasta \
    --kmer-length 16 \
    --output quality_control
```

With output:

```
Get spectra-cn plots and QV stats
meryl k=16 count output quality_control/merqury/FWD.maryl decontamination/decontaminated_fwd.fastq && meryl k=16 count output quality_control/merqury/
ntrol/merqury/UNION.maryl quality_control/merqury/FWD.maryl quality_control/merqury/REV.maryl && $MERQURY/merqury.sh quality_control/merqury/UNION.mar
INFO: With outputs {'merqury_output_dir': ''}
task completed successfully
        checkm_report_table : quality_control/checkM_Lineage/report_table.txt
        quast_report_html : quality_control/quast/quast/report.html
        merqury_output_dir : quality_control/merqury/
check quality_control for other reports

real    0m44.953s
user    6m32.068s
sys     0m18.196s
(base) wgalp:~/shared/144_working_directory>
```

This is the Quast plot of cumulative node length:

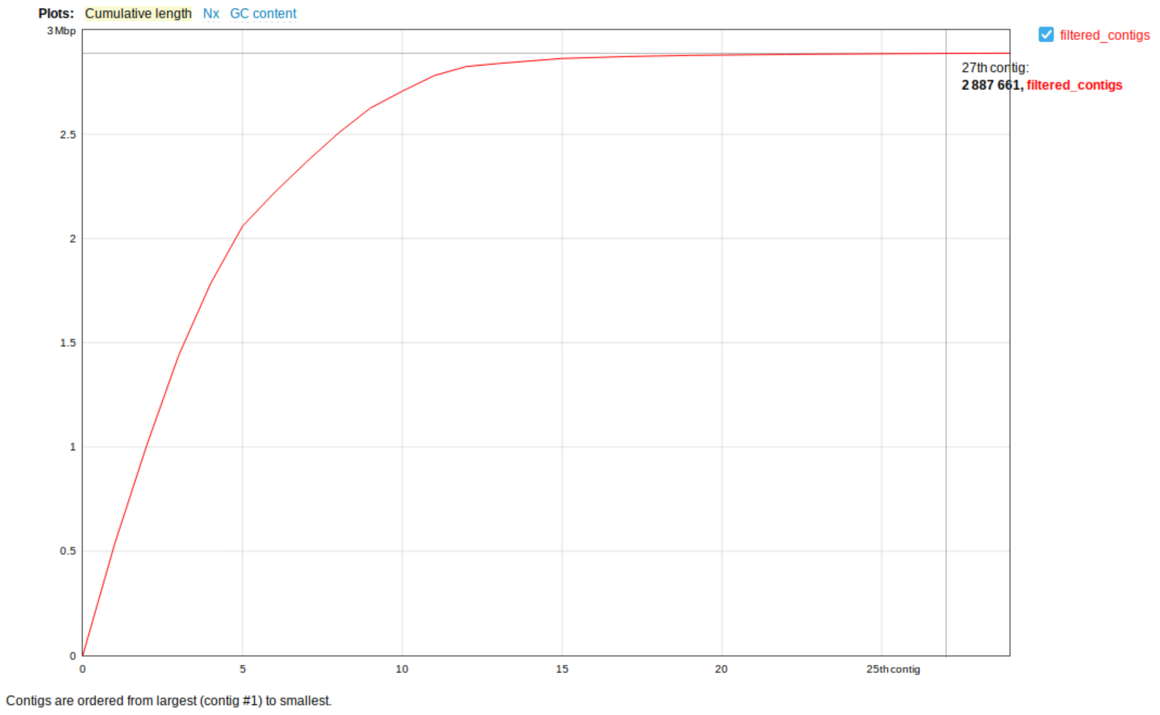

With Merqury, it is possible to see kmer multiplicity distribution:

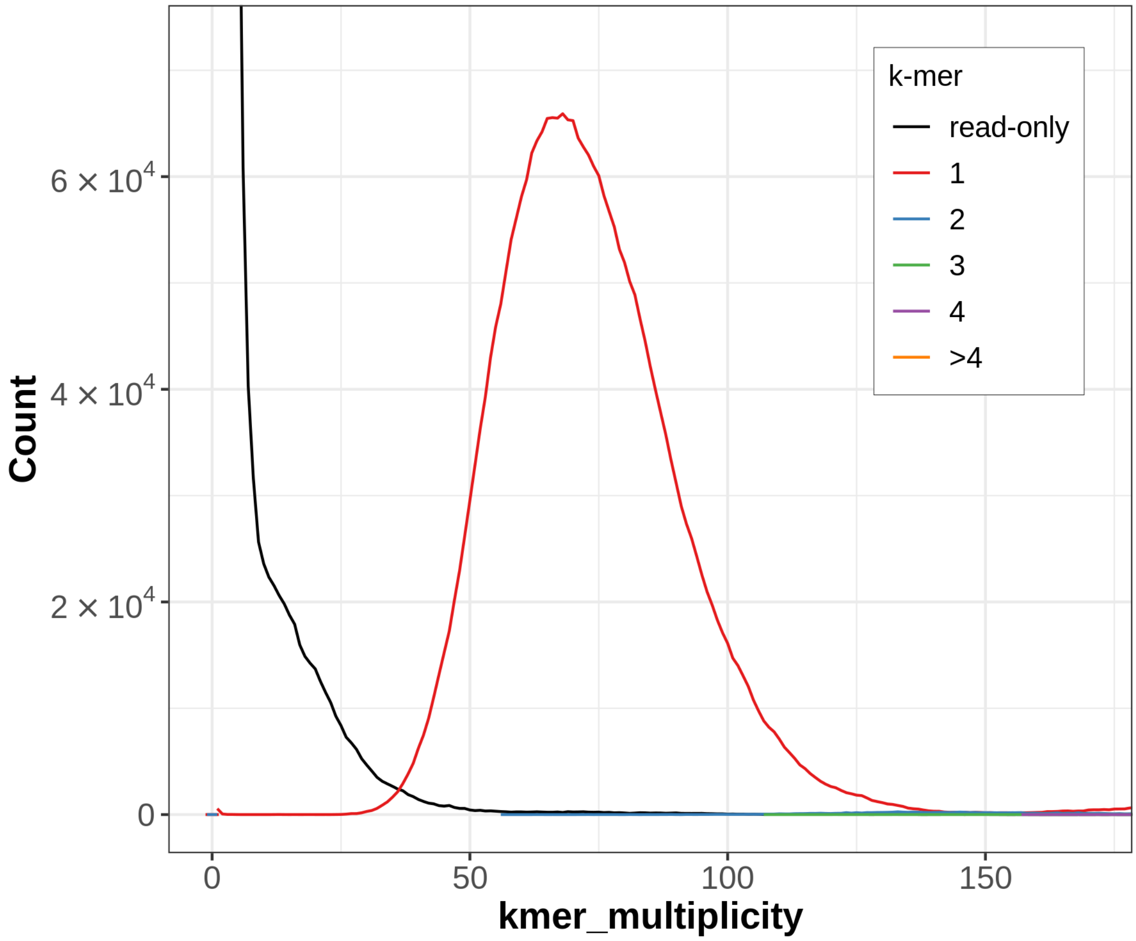

and the distribution here is that expected from an haploid genome.

CheckM can, among other things, compute tables of possible contamination of genomes. The output produced with `wgalp quality` is that of `lineage_wf` mode:

> CheckM requires ~15GB of available RAM to run in `linaeage_wf` mode with the *reduced tree* option

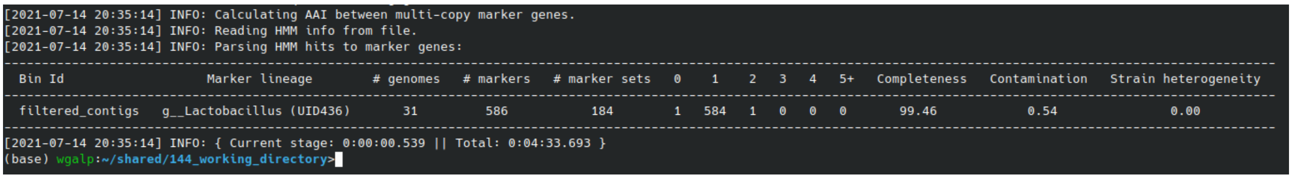

```
[2021-07-14 20:35:14] INFO: Calculating AAI between multi-copy marker genes.
[2021-07-14 20:35:14] INFO: Reading HMM info from file.
[2021-07-14 20:35:14] INFO: Parsing HMM hits to marker genes:
```

| Bin Id | Marker lineage | # genomes | # markers | # marker sets | 0 | 1 | 2 | 3 | 4 | 5+ | Completeness | Contamination | Strain heterogeneity |
|---|---|---|---|---|---|---|---|---|---|---|---|---|---|
| filtered_contigs | g__Lactobacillus (UID436) | 31 | 586 | 184 | 1 | 584 | 1 | 0 | 0 | 0 | 99.46 | 0.54 | 0.00 |

```
[2021-07-14 20:35:14] INFO: { Current stage: 0:00:00.539 || Total: 0:04:33.693 }
(base) wgalp:~/shared/144_working_directory>
```

To use the `taxonomic_wf`, run checkM manually:

```
# checkm taxonomy_wf -x <extension_of_taxa_file_to_use> \
#   <Rank> '<Taxon>' \
#   <input_folder> <output_folder>
# --- example ---
checkm taxonomy_wf -x .fasta \
    species 'Lactobacillus rhamnosus' \
    . taxa_checkm
```

It is possible to check which Ranks an Taxons are available in CheckM with the command:

```
checkm taxon_list | less
```

> while visualizing files with `less`, use the arrows or page up/down buttons to move inside the document. Press q to exit.

CheckM main output is discussed in the section *Comparison with shovill pipeline*.

##### NCBI compliant annotation using Prokka

If the user wants to deposit his/her genomes, he/she is required to annotate them. To this end, WGA-LP includes an interface to **Prokka** annotator that helps in creating NCBI compliant annotations:

```
(base) wgalp:~/shared/144_working_directory>wgalp annotate
--- no arguments given, printing help message ---
Wrapper to easily run prokka in NCBI compliant mode
--- arguments:
        --contigs : assembled contigs or scaffolds (.fasta)
        --output : path to the output folder
        --help : print this message

(base) wgalp:~/shared/144_working_directory>
```

In our example:

```
wgalp annotate \
    --contigs reordering/mauve_reorder/alignment2/filtered_contigs.fasta \
    --output annotation
```

```
[13:53:43] Thank you, come again.
INFO: With outputs {'ffn': 'prokka_annotated_genome.fnn', 'faa': 'prokka_annotated_genome.faa', 'gbk': 'prokka_annotated_genome.gbk', 'gff': 'prokka_annotated_genome.gff', 'tsv': 'prokka_annotated_genome.tsv'}
task completed successfully
the annotated assembly is at the following locations:
        FORMAT  PATH
        ffn     annotation/prokka/prokka_annotated_genome.fnn
        faa     annotation/prokka/prokka_annotated_genome.faa
        gbk     annotation/prokka/prokka_annotated_genome.gbk
        gff     annotation/prokka/prokka_annotated_genome.gff
        tsv     annotation/prokka/prokka_annotated_genome.tsv
other formats are available in the output folder annotation

real    1m57.486s
user    9m2.445s
sys     0m38.592s
(base) wgalp:~/shared/144_working_directory>
```

The produced annotated files can then be used for any downstream analysis.

This is the last step of the standard workflow for WGA-LP.

#### Other procedures

##### Load and unload the Kraken2 database into a RAMDisk

If it is planned to kraken2, or bracken multiple times, it may be useful to load the kraken2 database directly in RAM. To do so, use the `wgalp kdb-load` and `wgalp kdb-unload` procedures:

```
(base) wgalp:~/shared/144_working_directory>wgalp kdb-load
--- no arguments given, printing help message ---
Load kraken-db into RAM, this step is useful when running kraken multiple times.
Remember to unload the db after use and to use the --memory-mapped flag when supported (otherwise another copy of the db will be loaded).
--- arguments:
        --db : path to kraken database
        --mount-point : path where the kraken_db will be mounted
        --help : print this message

(base) wgalp:~/shared/144_working_directory>wgalp kdb-unload
--- no arguments given, printing help message ---
Unload kraken-db from RAM
--- arguments:
        --mount-point : path where the kraken_db was mounted
        --help : print this message

(base) wgalp:~/shared/144_working_directory>
```

For example like this:

```
wgalp:~/shared/144_working_directory>wgalp kdb-load --db $kraken_db --mount-point kraken_ramdisk
force True
WARNING: erasing an old output directory
INFO: Step kraken_ramdisk
INFO: Running on {'kraken_db': '/root/kraken_db'}
INFO:running >> mount -t tmpfs -o size=8G tmpfs kraken_ramdisk/kraken_ramdisk && cp -R /root/kraken_db kraken_ramdisk/kraken_ramdisk/
kraken_db
INFO: With outputs {'kraken_ram_db': 'kraken_db', 'kraken_ramdisk': ''}
task completed successfully
Kraken2 database loaded:
        database location (for --db arguments) : kraken_ramdisk/kraken_ramdisk/kraken_db
        ramdisk location (for wgalp kdb-unload) : kraken_ramdisk/kraken_ramdisk/
unload this ramdisk with the following command:
wgalp kdb-unload --mount-point kraken_ramdisk/kraken_ramdisk/
wgalp:~/shared/144_working_directory>wgalp kdb-unload --mount-point kraken_ramdisk/kraken_ramdisk/
task completed successfully
Kraken2 database unloaded
wgalp:~/shared/144_working_directory>
```

It could be useful to save the path to the loaded kraken2 database into a variable:

```
# use 'database location' path from wgalp kdb-load
kraken_ramdb=kraken_ramdisk/kraken_ramdisk/kraken_db
```

##### Filter FASTQ reads by ID

If needed, it is possible to filter fastq files by read ID, this can be useful to try different decontamination approaches, as seen in the next subsection:

```
(base) wgalp:~/shared/144_working_directory>wgalp filter-fastq
--- no arguments given, printing help message ---
Select reads from a fastq file by ID
--- arguments:
        --fastq : fastq reads to be filtered (.fastq)
        --fastq-fwd : fastq reads to be filtered FWD (Paired end mode) (.fastq)
        --fastq-rev : fastq reads to be filtered REV (Paired end mode) (.fastq)
        --selected-reads : a file containing the ids of the selected reads (each id is in its own line)
        --complement : if set, selected reads are instead removed from the original fastq
        --output : path to the output folder
        --help : print this message

(base) wgalp:~/shared/144_working_directory>
```

##### Use Kraken2 for decontamination

As an example, we show how it is possible to decontaminate raw fastq reads using kraken2 classification for reads.

> Even if interesting, we find this approach too aggressive, this is why we developed a different decontamination technique. This option is, however, of course easier and faster to run.

We consider the output (`kraken.log`) of kraken2 computed after decontamination (see the relative section), that can be simply obtained with `wgalp understand-origin`.

In our example, let us imagine that we want to keep only the reads that are recognized as originated from Lactobacillus rhamnosus. We can extract them using this command:

```
cat kraken_after_decontamination/kraken/kraken.log | \
    cut -d$'\t' -f 2,3 | \
    grep "Lactobacillus rhamnosus" | \
    cut -d$'\t' -f 1 > rhamnosus_reads.txt
```

This will generate `rhamnosus_reads.txt` (a text file with a read ID per line) that can be used with `wgalp filter-fastq`:

```
wgalp filter-fastq \
    --fastq-fwd decontamination/decontaminated_fwd.fastq \
    --fastq-rev decontamination/decontaminated_rev.fastq \
    --selected-reads rhamnosus_reads.txt \
    --output only_rhamnosus_reads
```

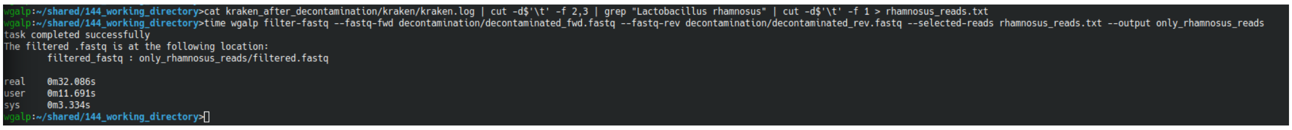

The output folder will contain the filtered reverse and forward reads.

#### Comparison with current state-of-the-art

In this section we show how the filtering used by our pipeline can improve the resulting Whole Genome Assembly. In particular, we compare the completeness and contamination metrics of **checkM** by computing the assembled genome with four approaches:

- By running a *blind* analysis with the **shovill** pipeline, that includes no decontamination step.
- By using **kraken2** classification for decontamination.
- By following the complete workflow of **WGA-LP**.
- By executing **ProDeGe** software for the decontamination of the final assembly

> With *blind* analysis, we mean that we do not apply any method to filter neither of the input data or the results

To run the **shovill** pipeline, we used the following command:

```
shovill \
    --R1 ../144/144_S13_L001_R1_001.fastq \
    --R2 ../144/144_S13_L001_R2_001.fastq \
    --tmpdir temp \
    --outdir out \
    --trim
```

The resulting assembly has the following checkM metrics:

```
(base) wgalp:~/shared/shovill_144/out>checkm taxonomy_wf -x .fasta species 'Lactobacillus rhamnosus' . taxa_checkm
[2021-07-14 06:31:41] INFO: CheckM v1.1.3
[2021-07-14 06:31:41] INFO: checkm taxonomy_wf -x .fasta species Lactobacillus rhamnosus . taxa_checkm
[2021-07-14 06:31:41] INFO: [CheckM - taxon_set] Generate taxonomic-specific marker set.
[2021-07-14 06:31:49] INFO: Marker set for Lactobacillus rhamnosus contains 952 marker genes arranged in 246 sets.
[2021-07-14 06:31:49] INFO: Marker set inferred from 10 reference genomes.
[2021-07-14 06:31:49] INFO: Marker set for Lactobacillus contains 409 marker genes arranged in 155 sets.
[2021-07-14 06:31:49] INFO: Marker set inferred from 135 reference genomes.
[2021-07-14 06:31:49] INFO: Marker set for Lactobacillaceae contains 396 marker genes arranged in 153 sets.
[2021-07-14 06:31:49] INFO: Marker set inferred from 143 reference genomes.
[2021-07-14 06:31:49] INFO: Marker set for Lactobacillales contains 335 marker genes arranged in 183 sets.
[2021-07-14 06:31:49] INFO: Marker set inferred from 490 reference genomes.
[2021-07-14 06:31:49] INFO: Marker set for Bacilli contains 250 marker genes arranged in 136 sets.
[2021-07-14 06:31:49] INFO: Marker set inferred from 821 reference genomes.
[2021-07-14 06:31:49] INFO: Marker set for Firmicutes contains 172 marker genes arranged in 99 sets.
[2021-07-14 06:31:49] INFO: Marker set inferred from 1349 reference genomes.
[2021-07-14 06:31:49] INFO: Marker set for Bacteria contains 104 marker genes arranged in 58 sets.
[2021-07-14 06:31:49] INFO: Marker set inferred from 5449 reference genomes.
[2021-07-14 06:31:49] INFO: Marker set written to: taxa_checkm/Lactobacillus rhamnosus.ms
[2021-07-14 06:31:49] INFO: { Current stage: 0:00:07.482 || Total: 0:00:07.482 }
[2021-07-14 06:31:49] INFO: [CheckM - analyze] Identifying marker genes in bins.
[2021-07-14 06:31:49] INFO: Identifying marker genes in 1 bins with 1 threads:
    Finished processing 1 of 1 (100.00%) bins.
[2021-07-14 06:35:43] INFO: Saving HMM info to file.
[2021-07-14 06:35:43] INFO: { Current stage: 0:03:54.021 || Total: 0:04:01.503 }
[2021-07-14 06:35:43] INFO: Parsing HMM hits to marker genes:
    Finished parsing hits for 1 of 1 (100.00%) bins.
[2021-07-14 06:35:44] INFO: Aligning marker genes with multiple hits in a single bin:
    Finished processing 1 of 1 (100.00%) bins.
[2021-07-14 06:36:39] INFO: { Current stage: 0:00:55.732 || Total: 0:04:57.236 }
[2021-07-14 06:36:39] INFO: Calculating genome statistics for 1 bins with 1 threads:
    Finished processing 1 of 1 (100.00%) bins.
[2021-07-14 06:36:39] INFO: { Current stage: 0:00:00.358 || Total: 0:04:57.595 }
[2021-07-14 06:36:39] INFO: [CheckM - qa] Tabulating genome statistics.
[2021-07-14 06:36:39] INFO: Calculating AAI between multi-copy marker genes.
[2021-07-14 06:36:39] INFO: Reading HMM info from file.
[2021-07-14 06:36:39] INFO: Parsing HMM hits to marker genes:
    Finished parsing hits for 1 of 1 (100.00%) bins.
-----------------------------------------------------------------------------------------------------------------------------------------------
  Bin Id          Marker lineage          # genomes   # markers   # marker sets   0     1     2     3    4    5+   Completeness   Contamination   Strain heterogeneity
-----------------------------------------------------------------------------------------------------------------------------------------------
  spades    Lactobacillus rhamnosus (6)       10          952           246         7    219   582   130   8    6       98.52           86.54                0.93
-----------------------------------------------------------------------------------------------------------------------------------------------
[2021-07-14 06:36:40] INFO: { Current stage: 0:00:01.267 || Total: 0:04:58.862 }
(base) wgalp:~/shared/shovill_144/out>
```

Using **kraken2** only:

```
[2021-07-18 16:50:46] INFO: Reading HMM info from file.
[2021-07-18 16:50:46] INFO: Parsing HMM hits to marker genes:
    Finished parsing hits for 1 of 1 (100.00%) bins.
-----------------------------------------------------------------------------------------------------------------------------------------------
  Bin Id            Marker lineage          # genomes   # markers   # marker sets   0     1    2   3   4   5+   Completeness   Contamination   Strain heterogeneity
-----------------------------------------------------------------------------------------------------------------------------------------------
  scaffolds    Lactobacillus rhamnosus (6)     10          952           246         18    930   3   0   1   0       97.75           0.38                0.00
-----------------------------------------------------------------------------------------------------------------------------------------------
[2021-07-18 16:50:47] INFO: { Current stage: 0:00:00.554 || Total: 0:02:27.894 }
wgalp:~/shared/144_working_directory/only_kraken2_test/assembly/SPAdes/scaffolds>
```

With **WGA-LP** pipeline:

```
[2021-07-14 06:55:23] INFO: { Current stage: 0:00:01.152 || Total: 0:02:38.676 }
[2021-07-14 06:55:23] INFO: Calculating genome statistics for 1 bins with 1 threads:
    Finished processing 1 of 1 (100.00%) bins.
[2021-07-14 06:55:23] INFO: { Current stage: 0:00:00.219 || Total: 0:02:38.895 }
[2021-07-14 06:55:23] INFO: [CheckM - qa] Tabulating genome statistics.
[2021-07-14 06:55:23] INFO: Calculating AAI between multi-copy marker genes.
[2021-07-14 06:55:23] INFO: Reading HMM info from file.
[2021-07-14 06:55:23] INFO: Parsing HMM hits to marker genes:
    Finished parsing hits for 1 of 1 (100.00%) bins.
-----------------------------------------------------------------------------------------------------------------------------------------------
  Bin Id                Marker lineage          # genomes   # markers   # marker sets   0     1    2   3   4   5+   Completeness   Contamination   Strain heterogeneity
-----------------------------------------------------------------------------------------------------------------------------------------------
  filtered_contigs   Lactobacillus rhamnosus (6)     10          952          246         11   934   6   0   1   0       98.19          1.27                0.00
-----------------------------------------------------------------------------------------------------------------------------------------------
[2021-07-14 06:55:24] INFO: { Current stage: 0:00:00.908 || Total: 0:02:39.804 }
(base) wgalp:~/shared/144_working_directory>
```

To test **ProDeGe** pipeline for automatic contig filtering we launched the program using the assembly obtained from the trimmed reads:

```
[2021-07-25 20:36:46] INFO: [CheckM - qa] Tabulating genome statistics.
[2021-07-25 20:36:46] INFO: Calculating AAI between multi-copy marker genes.
[2021-07-25 20:36:46] INFO: Reading HMM info from file.
[2021-07-25 20:36:46] INFO: Parsing HMM hits to marker genes:
    Finished parsing hits for 1 of 1 (100.00%) bins.
-----------------------------------------------------------------------------------------------------------------------------------------------
  Bin Id                   Marker lineage          # genomes   # markers   # marker sets   0     1     2    3    4   5+   Completeness   Contamination   Strain heterogeneity
-----------------------------------------------------------------------------------------------------------------------------------------------
  LacRhN132_output_clean   Lactobacillus rhamnosus (6)     10          952          246         7    235   662   40   6   2       98.52          72.99               1.19
-----------------------------------------------------------------------------------------------------------------------------------------------
[2021-07-25 20:36:47] INFO: { Current stage: 0:00:01.137 || Total: 0:05:21.560 }
(base) redsnic@redsnic-MS-7760:~/WGA_batteri/LacRhN132_output_clean$
```

As we will show in the final table, ProDeGe retains many nodes that are assembled from Pedicoccus reads. Removing the pure Pediococcus scaffolds with kraken2 in combination with `wgalp filter-assembly` widely improves the assembly, producing metrics similar to that of WGA-LP.

The following table summarizes some relevant specifications on the assembled genomes:

| Feature | shovill | Kraken2 | WGA-LP | ProDeGe | refined ProDeGe |
|---|---|---|---|---|---|
| **GC** | 0.44308 | 0.46934 | 0.46714 | 0.44758 | 0.46711 |
| **GC std** | 0.06668 | 0.02220 | 0.01823 | 0.02804 | 0.00932 |
| **Genome size** | 5262721 | 2733655 | 2892519 | 4894038 | 2871414 |
| **# ambiguos bases** | 0 | 300 | 100 | 210 | 200 |
| **# scaffolds** | 453 | 83 | 40 | 34 | 15 |
| **Longest scaffold** | 535921 | 412497 | 535910 | 535910 | 535910 |
| **N50** | 222429 | 142270 | 345213 | 195725 | 431489 |
| **Mean scaffold length** | 11617.486 | 32935.602 | 72312.975 | 143942.29 | 191427.6 |
| **coding density** | 0.85661 | 0.84790 | 0.85164 | 0.85936 | 0.85315 |
| **# predicted genes** | 5337 | 2618 | 2751 | 4673 | 2714 |

At the price of a very small (possible) loss of completeness, the contamination is drastically reduced by using Kraken2 or WGA-LP. The table shows how WGA-LP decontamination is less strict than kraken2 selection in eliminating reads, reducing the probability of discarding reads from the target organism.

From NCBI's Lactobacillus Rhamnosus web page we can get the following table:

| Feature | Value |
|---|---|
| **median total length (Mb)** | 2.949 |
| **median protein count** | 2652 |
| **median GC%** | 46.7 |

That is colse to the results we achieved with WGA-LP.

> The goal of this comparison is to show how important is to take care of the details when doing whole genome assemblies

##### Reproducing this analysis

Finally, we have shown with an example that WGA-LP can produce high quality Whole Genome Assemblies even with contaminated data.

To reproduce the analysis as seen in this document the following steps have to be performed:

- **Installation**: use `Docker` installation
- **Raw Reads**: download the reads from **SRA** (with SRA ID **SRR15265000**, BioProject **PRJNA749304**)
- **Directory setup**: Create folders named `144`, `144_working_directory` and ‘`references` with the reads in the `/root/shared` directory

- **References**: For decontamination, the accession numbers of the references are reported in the the following table. Save the references in `/root/shared/references/subfolder` according to the following table

| Organism | Subfolder name | Accession Numbers |
|---|---|---|
| **Lactobacillus rhamnosus** | rhamnosus | `NZ_CP040780.1, NZ_CP021426.1, NC_017491.1, NZ_CP067042.1, NZ_CP014201.1, NZ_CP046267.1, NZ_CP044506.1, NZ_LT220504.1, NZ_CP073317.1, NZ_CP006804.1, NZ_CP031290.1, NC_017482.1, NC_013198.1, NZ_CP046395.1, NZ_CP022109.1, NZ_CP067365.1, NC_021723.1, NC_021725.1, NZ_CP017063.1, CP016823.1, NZ_CP025428.1, NZ_CP053619.1, NC_013199.1, NZ_LR698954.1, NZ_LR134322.1, NZ_LR134331.1, NZ_CP020464.1, NZ_CP019305.1, NZ_CP045586.1, NZ_CP073711.1, NZ_CP044228.1` |

| Organism | Subfolder name | Accession Numbers |
|---|---|---|
| **Pediococcus Acidilactici** | pediococcus | NZ_CP033438.1, NZ_CP018763.1, NZ_CP048019.1, NZ_CP066046.1, NZ_CP066066.1, NZ_CP068106.1, NZ_CP061715.1, NZ_CP023654.1, NZ_CP025471.1, CP050079.1, NZ_CP053421.1, CP021487.1, NZ_CP021484.1, NZ_CP021529.1, NZ_CP028247.1, NZ_CP028249.1, NZ_CP035154.1, NZ_CP035266.1, NZ_CP015206.1, NZ_CP067392.1, NZ_CP035151.1 |

The commands are run from the `144_working_directory`, you can check the exact location in the images of this manual.
